# BLADE-ON-PETIOLE interacts with CYCLOIDEA to fine-tune *CYCLOIDEA*-mediated flower symmetry in Monkeyflowers (*Mimulus*)

**DOI:** 10.1101/2024.01.08.574647

**Authors:** Yuan Gao, Jingjian Li, Jiayue He, Yaqi Yu, Zexin Qian, Zhiqiang Geng, Liuhui Yang, Yumin Zhang, Yujie Ke, Qiaoshan Lin, Jing Wang, Sumei Chen, Fadi Chen, Yao-Wu Yuan, Baoqing Ding

## Abstract

Morphological novelties, or key innovations, are instrumental to the diversification of the organisms. In plants, one such innovation is the evolution of zygomorphic flowers, which is thought to promote outcrossing and increases flower morphological diversity. We isolated three allelic mutants from two *Mimulus* species displaying altered floral symmetry and identified the causal gene as the orthologue of *Arabidopsis BLADE-ON-PETIOLE*. We found that MlBOP and MlCYC2a physically interact and this BOP-CYC interaction module is highly conserved across the angiosperms. Furthermore, MlBOP self-ubiquitinates and suppresses *MlCYC2a* self- activation. MlCYC2a, in turn, impedes MlBOP ubiquitination. Thus, this molecular tug-of-war between MlBOP and MlCYC2a fine-tunes the expression of *MlCYC2a*, contributing to the formation of bilateral symmetry flowers, a key trait in angiosperm evolution.

**One Sentence Summary:** Molecular tug-of-war between MlBOP and MlCYC2a fine-tunes the expression of *MlCYC2a*, contributing to the bilateral flower symmetry formation.

## Main Text

Morphological novelties, or key innovations, are instrumental to the diversification of the organisms (*1, 2*). In plants, one such innovation is the evolution of zygomorphic flowers, which differentiate along the dorsiventral axis, and is thought to enhance diverse pollinator exploitation for promoting outcrossing (*3*) and increases flower morphological diversity (*2*). Floral zygomorphy has evolved multiple times independently in many angiosperm lineages from their actinomorphic ancestors (*4–6*). Due to its prevalence, essential ecological functions and evolutionary implications, the molecular mechanisms underpinning floral zygomorphy have been extensively investigated in many plant lineages. The common garden snapdragon (*Antirrhinum*) is one of the best investigated systems: two functionally redundant and dorsally expressed genes, *CYCLOIDEA* (*CYC*) and *DICHOTOMA* (*DICH*) (*7, 8*), activate *RADIALIS* (*RAD*) (*9*) in the dorsal petals. *RAD*, in turn, competes with two DIV-RAD interacting factors (DRIFs) (*10*) to bind with *DIVARICATA* (*DIV*) (*11, 12*), establishing zygomorphy. This model has been widely adopted to explain the alteration of floral symmetry in many plant lineages.

Central to the model is the spatially restricted expression pattern of *CYC*, as expanding its expression leads to dorsalized actinomorphy (*7*). This spatial expansion of the *CYC*-like genes has been documented in various taxa, converting flowers from zygomorphy to actinomorphy (*13–22*). However, the evolutionary shifts from zygomorphy to actinomorphy have often involved a reduced number of petals, unlike the increased number observed in the *Antirrhinum cyc* mutant (*11*), implying additional underlying mechanisms beyond simple changes in the *CYC* expression domain (*4, 23, 24*). Despite that the association between the alteration of flower symmetry and the changes in *CYC* expression pattern has long been established, how the expression of *CYC* is regulated and whether additional genetic factors are involved remain largely unknown.

In this study, we employed a chemical mutagenesis approach to screen for mutants with altered flower symmetry in the monkeyflower species *Mimulus lewisii* and *M. verbenaceus*. We isolated three allelic mutants displaying altered floral symmetry, with significantly upregulated expression of *CYC*. Genomic and Sanger sequencing revealed three independent mutations in the orthologue of *Arabidopsis BLADE-ON-PETIOLE* (*BOP*) in these mutants. Functional analyses further validated that *BOP* is the causal gene underlying the mutants. The overlapping of the spatiotemporal expression patterns of *MlBOP* and *MlCYC2a* as revealed by *in situ* hybridization analysis suggests that *MlBOP* might directly regulate the expression of *MlCYC2a*.

Surprisingly, we found that MlBOP and MlCYC2a physically interact and this BOP- CYC interaction module is highly conserved across the angiosperms. Furthermore, we revealed that MlBOP functions as an E3 ligase adaptor to self-ubiquitinate, but does not ubiquitinate its interacting partner MlCYC2a. Instead, MlBOP suppresses the self- activation of *MlCYC2a*. MlCYC2a, on the other hand, impedes the ubiquitination of MlBOP. Thus, this molecular tug-of-war between MlBOP and MlCYC2a fine-tunes the expression of *MlCYC2a*, contributing to the formation of bilateral symmetry flowers in *Mimulus*.

## Results

### Phenotypic characterization of the floral symmetry mutants in *Mimulus*

The corolla of the wild-type *M. lewisii* (LF10) is composed of three types of petals arranged asymmetrically along the adaxial-abaxial axis: two larger dorsal petals on the adaxial side, two lateral petals, and one ventral petal on the abaxial side (Fig. 1A). Despite the well-established relationship between flower symmetry and the spatiotemporal expression pattern of *CYC* in many flowering plants (*25*), how the expression pattern of *CYC* is regulated remains elusive. To address this, we conducted a forward genetics screen for ethyl methanesulfonate (EMS) induced mutants displaying altered flower symmetry in the LF10 background. We isolated two morphologically similar mutants (ML14181 and ML14132). Both mutants exhibited a transition of flower symmetry from zygomorphy to actinomorphy, as the size of their lateral and ventral petals in both mutants was similar to the two dorsal petals (Fig. 1, B and C). Notably, unlike the ML14132 mutant, the ML14181 mutant flowers had four petals instead of five, along with misregulations of floral organ identity, such as the outgrowth of extra petaloid floral organs between the sepal and petal whorl, and the homeotic transformations in the stamen whorl (Fig. 1, D to F). Furthermore, the disruption of intra-organ boundaries was evident with the development of green tissues between the petals in both mutants (Fig. 1, E and F). The number of extra tissues also varied among flowers (table S1). To ascertain if the two phenotypically similar mutants were caused by the same gene, we conducted a complementation cross and found that the resulting F_1_ progeny failed to rescue the mutant phenotypes, indicating allelism. Thus, we focused on the more severe ML14181 mutant and characterized its floral morphology using scanning electron microscopy (SEM). Early flower development in the ML14181 mutant (Fig. 1, K to N) showed significant differences compared to the wild type (Fig. 1, G to J), including the presence of only four petal primordia and delayed or aborted stamen initiation (Fig. 1, K to M).

**Fig. 1.**
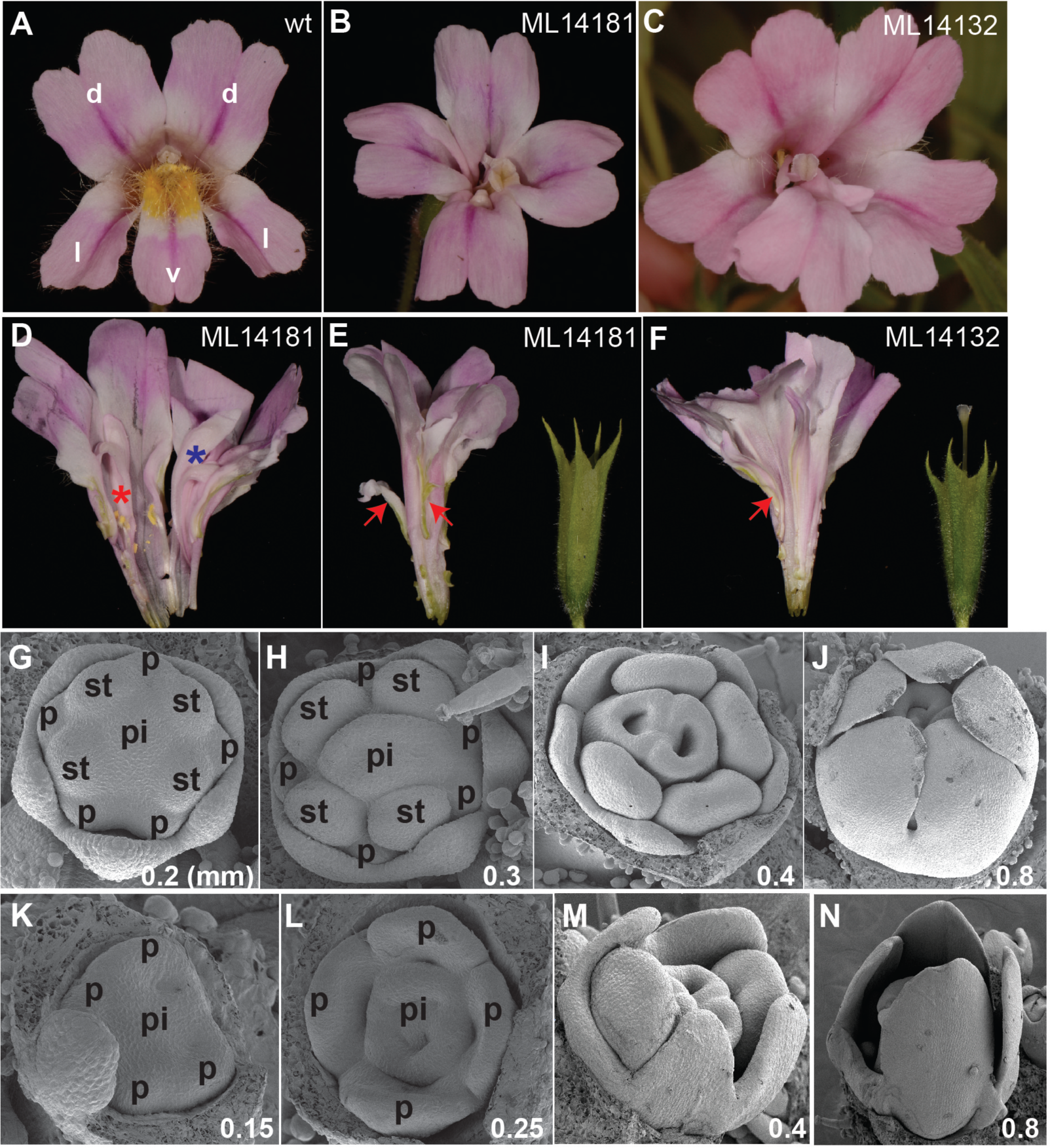
Phenotypic characterization of the two floral symmetry mutants in *Mimulus lewisii*. (**A, B, C**) Face view of the wild-type *M. lewisii* (inbred line LF10), ML14181 and ML14132 mutant corolla. d: dorsal; l: lateral; v: ventral. (**D**) Homeotic transitions from stamens to petaloid organs in ML14181. Red asterisk labels a petaloid organ, and blue asterisk labels an extra petaloid organ. (**E** and **F**) Side view of the ML14181 and ML14132 corolla and calyx. Arrows point to the extra tissues developed between the calyx and corolla whorl and the green tissues developed in the interprimordial region. (**G**-**J**) SEM on wt flower buds at different developmental stages. p: petal; st: stamens; pi: pistil. (**K**-**N**) SEM on the ML14181 flower buds at different developmental stages.

Additionally, we identified a new mutant, NJ01339, in *M. verbenaceus*, displaying nearly identical floral phenotypes to the two mutants in LF10. The wild-type *M. verbenaceus* corolla exhibited a similar symmetry plan to *M. lewisii* (Fig. 2A), while the NJ01339 mutant flowers had four to five nearly identical petals (Fig. 2B). To investigate if the mutations in the three mutants from the two species were allelic, we performed interspecies complementation crosses. The F_1_ hybrids derived from the mutants exhibited nearly radial symmetrical flowers with petaloid organs developed in the stamen whorl (Fig. 2, C to E). Interestingly, the F_1_ from the cross between ML14181 and NJ01339 (Fig. 2D) had more petaloid organs in the stamen whorl compared to the F_1_ from the cross between ML14132 and NJ01339 (Fig. 2E), suggesting that these three alleles were likely hypomorphic. Together, the results of the complementation crosses indicate that the mutants uncovered from *M. verbenaceus*, along with ML14181 and ML14132 mutants, likely represent three alleles of the same gene.

**Fig. 2.**
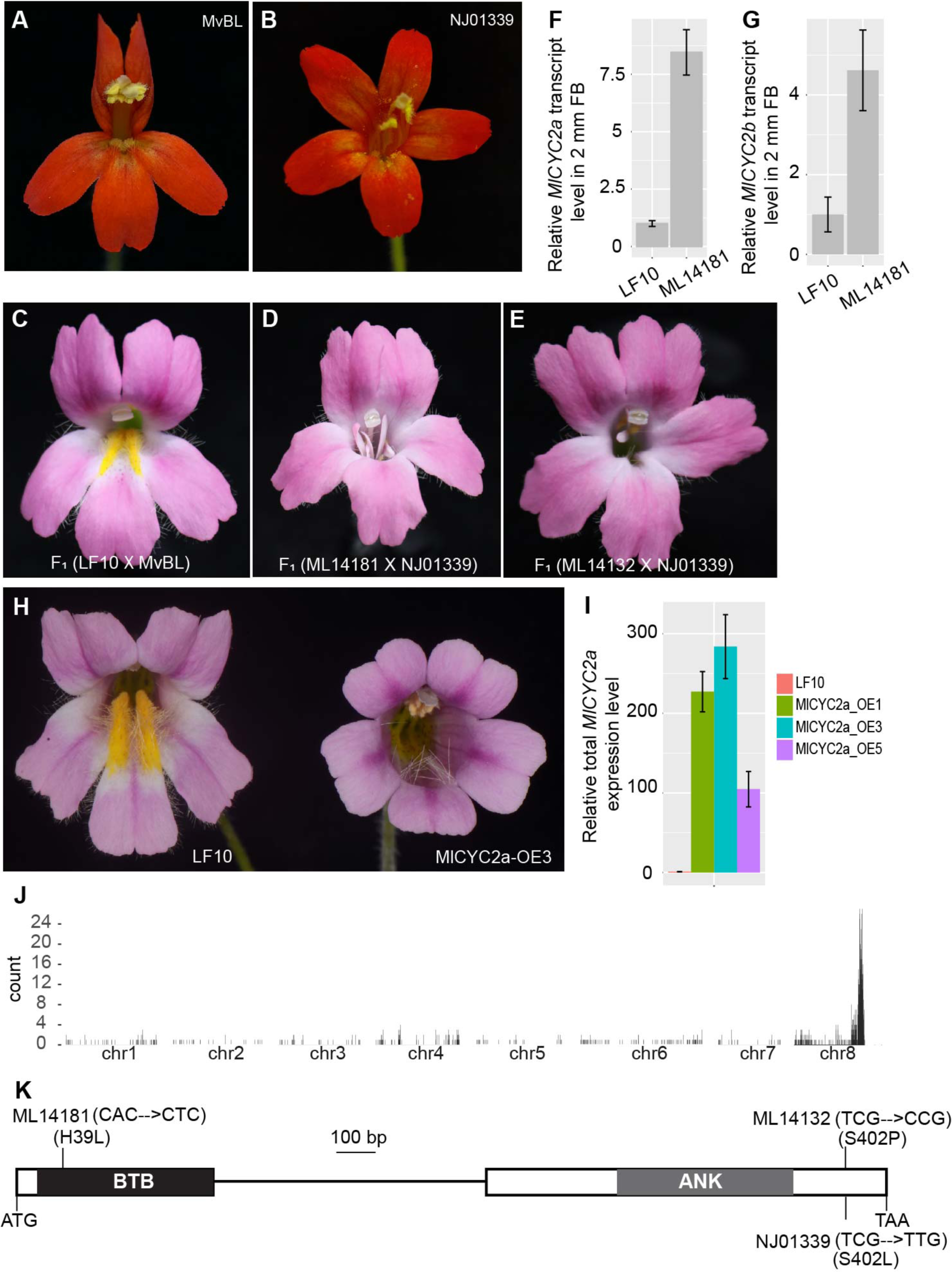
Identification of the gene responsible for the mutant flower phenotypes. (**A**) and (**B**) are the front view of the flower symmetry plan in *M.verbenaceus* and NJ01339, respectively. (**C**) Face view of the flower phenotype of the F_1_ derived from the interspecies hybridization between the wildtype *M.lewisii* and *M.verbenaceus*. (**D**-**E**) Flower phenotypes of the individuals derived from the interspecies complementation crosses between *M.lewisii* and *M.verbenaceus* mutants. (**F**-**G**) Quantitative measurement of *MlCYC2a* and *MlCYC2b* expression in wt and ML14181. *MlUBC* was used as the reference gene. Error bars represent 1 SD from three biological replicates. (**H**) Comparison of flower symmetry phenotype between wt and *35S*:*MlCYC2a* in the LF10 background. (**I**) Total *MlCYC2a* expression level in the wt and three representative *35S*:*MlCYC2a* transgenic lines. *MlUBC* was used as the reference gene. Error bars represent 1 SD from three biological replicates. (**J**) Genome scan for regions that are enriched in homozygous single nucleotide polymorphisms (SNP s) reveals a sharp peak (indicated by the asterisk). Each pseudoscaffold of the *Mimulus lewisii* SL 9 (the mapping line) genome was binned into 20-kb intervals, and the number of homozygous SNP s in each 20-kb interval was plotted in a bar graph. (**K**) Exon-intron structure of *Mimulus BOP.* A non-synonymous substitution (A to T) causes the amino acid replacement from Histidine to Leucine at position 39 (H39L) for ML14181 and a non-synonymous substitution (T to C) causes the amino acid replacement from Serine to Proline (S402P) for ML14132, and a non-synonymous substitution (C to T) causes the amino acid replacement from Serine to Leucine (S402L) for NJ01339. Black box: BTB domain; Grey box: Ankyrin repeats; White box: Coding region; Line: Intron. Scale bar is 100 bp.

### Shift from zygomorphy to actinomorphy in the mutants is correlated with the ectopic expression of *CYCLOIDEA* in *Mimulus*

Flower dorsalization is often associated with changes in the spatiotemporal expression of *CYC*-like genes (*3*). To investigate whether the expression of *CYC*-like genes is linked to the alteration of flower symmetry in the mutant flowers, we first identified the *CYC* orthologs in the LF10 genome. Similar to many species in Lamiales, we found two copies of the *CYC*-like genes named *MlCYC2a* and *MlCYC2b* (fig. S1A), following previous nomenclature schemes (*26*). Both genes were expressed at relatively low levels in LF10 flower buds across different developmental stages (fig. S1B). Notably, in the ML14181 background, *MlCYC2a* and *MlCYC2b* were upregulated by nearly 8-fold and 5-fold, respectively, as revealed by quantitative reverse transcription PCR assay (RT-qPCR) (Fig. 2, F and G), suggesting that the alteration of the *CYC* gene expression patterns may be linked to the shift from zygomorphy to actinomorphy in the mutants.

To determine whether the upregulation of *CYC-like* expression levels leads to the dorsalized flowers observed in the mutants, we chose *MlCYC2a* for overexpression using the cauliflower mosaic virus (CaMV) *35S* promoter in LF10 wild-type background, due to known functional redundancy of *CYC* paralogues in other Lamiales species (*26*). As expected, the transgenic lines with high *MlCYC2a* expression levels exhibited completely dorsalized flowers as seen in the three independent representative transgenic lines with similar floral phenotypes (Fig. 2, H and I). However, it is important to note that significant differences were observed between the floral phenotypes of the ML14181/ML14132 mutants and the *35S*: *MlCYC2a* transgenic lines. The mutants displayed extra floral organs and partial homeotic transformations, which were not observed in the *35S*: *MlCYC2a* transgenic lines, suggesting that the mutant likely regulates an array of additional genes related to flower development, in addition to *MlCYC2a* and *MlCYC2b*.

### Identification of the mutation in ML14181 as the ortholog of *Arabidopsis BLADE-ON-PETIOLE*

To identify the gene responsible for the ML14181 mutant phenotypes, we conducted a bulk segregant analysis and compared single-nucleotide polymorphism (SNP) profiles of several previously reported EMS mutants (table S2). This analysis revealed a single sharp peak at the end of the chromosome 8 (Fig. 2J), which corresponds to the gene encoding the *Arabidopsis BLADE-ON-PETIOLE* (*BOP*) homolog in *Mimulus* (fig. S2). BOP contains two conserved protein-protein interaction motifs: a BTB/POZ (for Broad Complex, Tramtrack, and Bric-a-brac/POX virus and Zinc finger) domain at the N-terminus and four ankryin motifs near the C-terminus (*27*). The identified gene in ML14181 contains a non-synonymous substitution, leading to a Histidine-to-Leucine replacement (H39L, Fig. 2K). Notably, this H residue at position 39 is highly conserved in all protein species containing a BTB/POZ domain across plants, animals, and fungi (*28, 29*). We reasoned that this H39L replacement is likely to disrupt protein function, making *BOP* the most promising candidate underlying the mutant phenotype.

Further analysis of the *BOP* coding sequence from ML14132 and NJ01339 mutants through Sanger sequencing revealed independent mutations. The ML14132 mutant carried a non-synonymous substitution of Serine-to-Proline (S402P, Fig. 2K) neighboring the ankryin repeat motif. Strikingly, NJ01339 had the exact same position being mutated as ML14132, but with a non-synonymous substitution of Serine-to-Leucine (S402L, Fig. 2K), further highlighting the importance of this residue. Taken together, the three independent mutations strongly support *BOP* as the causal gene responsible for the mutant phenotypes. Hence, we renamed the ML14181, ML14132, and NJ01339 mutants as *bop-1*, *bop-2* and *bop-3*, respectively. To further validate the function of *BOP*, we performed a CRISPR/Cas9-mediated knockout of the *BOP* in *M. verbenaceus.* The resulting frameshift mutations in *BOP* closely resembled the *bop-3* mutant phenotypes (Fig. 3, A and C). The two targeted mutations do not complement the *bop-3* mutant phenotypes, confirming that they are allelic (Fig. 3B).

**Fig. 3.**
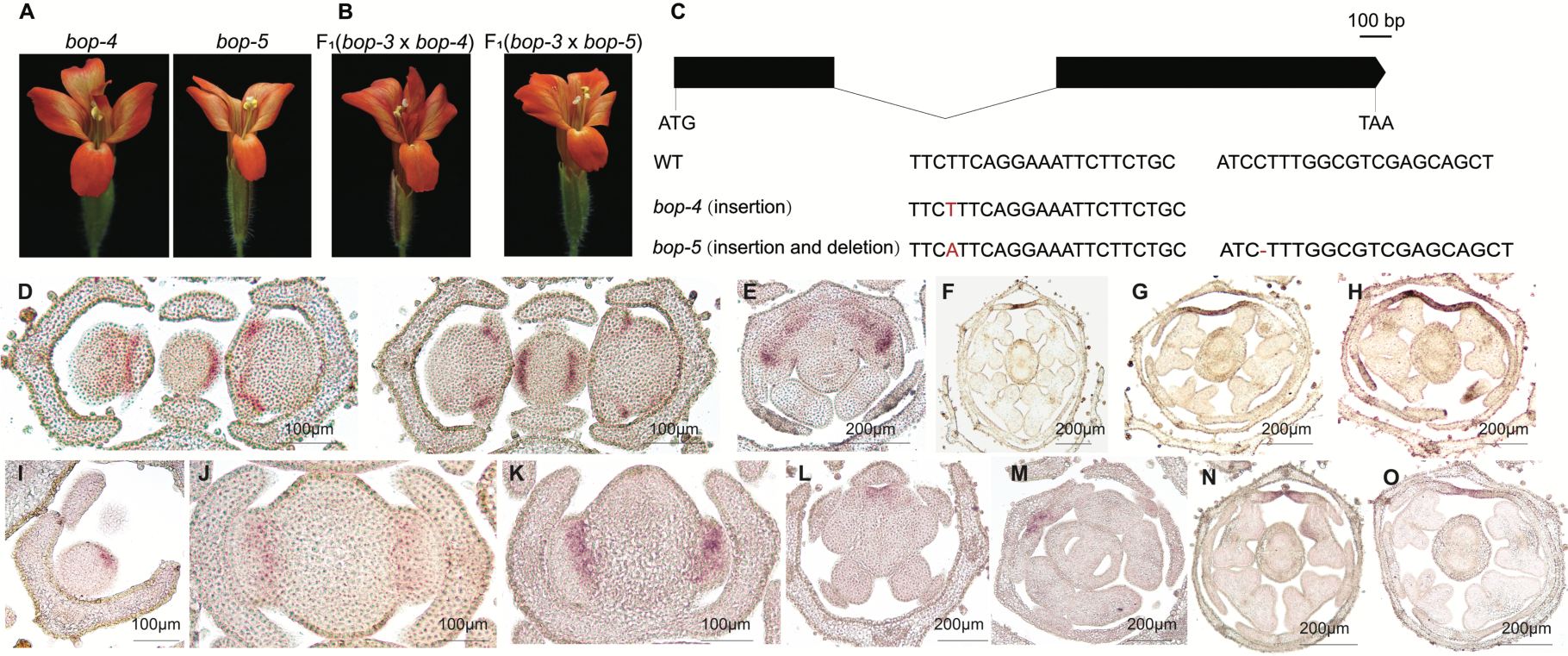
Functional characterization of the *BOP* in *M.verbenaceus* and *in situ* hybridization of *MlBOP* and *MlCYC2a* in the wildtype LF10 background. (**A**) Flower phenotypes of the two independent *BOP* CRISPR-Cas9 knockout lines. (**B**) Flower phenotypes of the complementation crosses between the two CRISPR lines and the *bop-3* mutant. (**C**) Illustrations of two independent CRISPR alleles relative to the guide RNA position in *MvBOP*. (**D**-**H**) Spatiotemporal expression pattern of *MlBOP* in LF10 flower buds at different developmental stages. (**I**-**O**) Spatiotemporal expression pattern of *MlCYC2a* in LF10 flower buds at different developmental stages.

To further confirm that *BOP* is the causal gene underlying the mutants, we attempted to complement the *bop-1* mutant with a wildtype *BOP* transgene. However, due to severe pistil development impairment in the mutant, we circumvented this by first overexpressing the *MlBOP* driven by *35S* promoter in the wild type background and then crossing this transgene into the *bop-1* mutant background. As expected, the transgene rescued the *bop-1* mutant phenotype, confirming that *BOP* is the causal gene. Similarly, introducing *35S*:*YFP-MlBOP* transgene into the *bop-3* background rescued the *bop-3* mutant phenotypes (fig. S3), further confirming that the observed mutant phenotypes are due to the loss of *BOP* function.

### Spatiotemporal expression of *MlBOP* overlaps with *MlCYC2a*

To explore the regulatory relationships between *MlBOP* and *MlCYC2a*, we traced the expression patterns of both genes in LF10 with *in situ* hybridization assay. *MlBOP* exhibited early expression at the junctions between the floral meristem and the shoot apical meristem (two consecutive sections in Fig. 3D). As the floral meristem develops, *MlBOP* expression became localized to the dorsal (adaxial) region of the floral meristem. Subsequently, *MlBOP* expression was spatially restricted to the two dorsal petals (Fig. 3, E to G). As flower buds further developed, *MlBOP* expression expanded to the lateral and ventral petals (Fig. 3H), which was independently validated by our RT-PCR analysis of the floral organs dissected from the wildtype 10 mm floral buds (fig. 1C).

Similarly, the expression of *MlCYC2a* was detected in a few cells at the adaxial side of the floral meristem (Fig. 3I). As the floral meristem expanded, *MlCYC2a* expression was restricted to the adaxial side of the floral primordia (Fig. 3, J and K). With the emergence of petal and stamen primordia, *MlCYC2a* expression remained restricted to the two adaxial petal and stamen primordia (Fig. 3L). As floral organs continued to differentiate, *MlCYC2a* expression was retained only in the two dorsal petal primordia throughout further development (Fig. 3, L to O), which was also independently validated by our RT-PCR analysis (fig. 1C). The expression pattern of *MlCYC2a* closely mirrored that of *MlBOP* at early developmental stages, suggesting that *MlBOP* might regulate *MlCYC2a*.

### MlBOP physically interacts with MlCYC2a and MlCulline3a

In Arabidopsis, AtBOP2 has been reported as an E3 ubiquitin ligase complex to regulate PHYTOCHROME INTERACTING FACTOR 4 and LEAFY to regulate plant development (*30, 31*). The overlapping expression domains between *MlBOP* and *MlCYC2a* prompted us to speculate that MlBOP may function as a putative member of a CUL3^BOP^ E3 ubiquitin ligase complex to post-translationally modify MlCYC2a. To test this hypothesis, we conducted yeast two-hybrid assays to examine the interactions between MlBOP, MlCYC2a, and MlCulline3a. The results indicated that MlBOP and MlCulline3a interacted weakly in yeast. Surprisingly, the interaction between bop-1 and MlCulline3a appeared stronger (Fig. 4A). We also found that bop- 1 forms homodimers in yeast, however, we were not able to test this for the wild-type MlBOP due to its self-activation in yeast cells. Additionally, we found that MlCYC2a and MlCYC2b can form homodimers and heterodimers as previously reported in other species (fig. S4) (*32*). Most importantly, we found that both MlBOP and bop-1 can interact with MlCYC2a and MlCYC2b, and bop-1 appears to interact stronger with MlCYC2a and MlCYC2b than MlBOP in yeast (Fig. 4A, fig. S4). The interaction between MlBOP, bop-1 and MlCYC2a was further confirmed in a pull-down assay. Consistently, bop-1 seems to be able to pull-down a larger quantity of MlCYC2a than MlBOP, suggesting that bop-1 interacts stronger with MlCYC2a than MlBOP (Fig. 4B). To further confirm these interactions in planta, we employed a bimolecular fluorescence complementation assay (BiFC) (Fig. 4C). However, we were not able to compare the affinities between the MlBOP-MlCYC2a and bop- MlCYC2a directly as the intensity of the fluorescence signal is difficult to compare across different experiments. Therefore, we utilized surface plasmon resonance (SPR) assay to measure the binding affinities (*K_D_*) of the MlBOP-MlCYC2a and bop-1- MlCYC2a complexes in real time. The SPR assay revealed that the *K_D_* of the bop-1- MlCYC2a complex was stronger (*K_D_* = 9.68E-7 M) than that of the MlBOP-MlCYC2a complex (*KD* = 1.44E-6 M) (Fig. 4D), which is consistent with our Y2H and pull-down results.

**Fig. 4.**
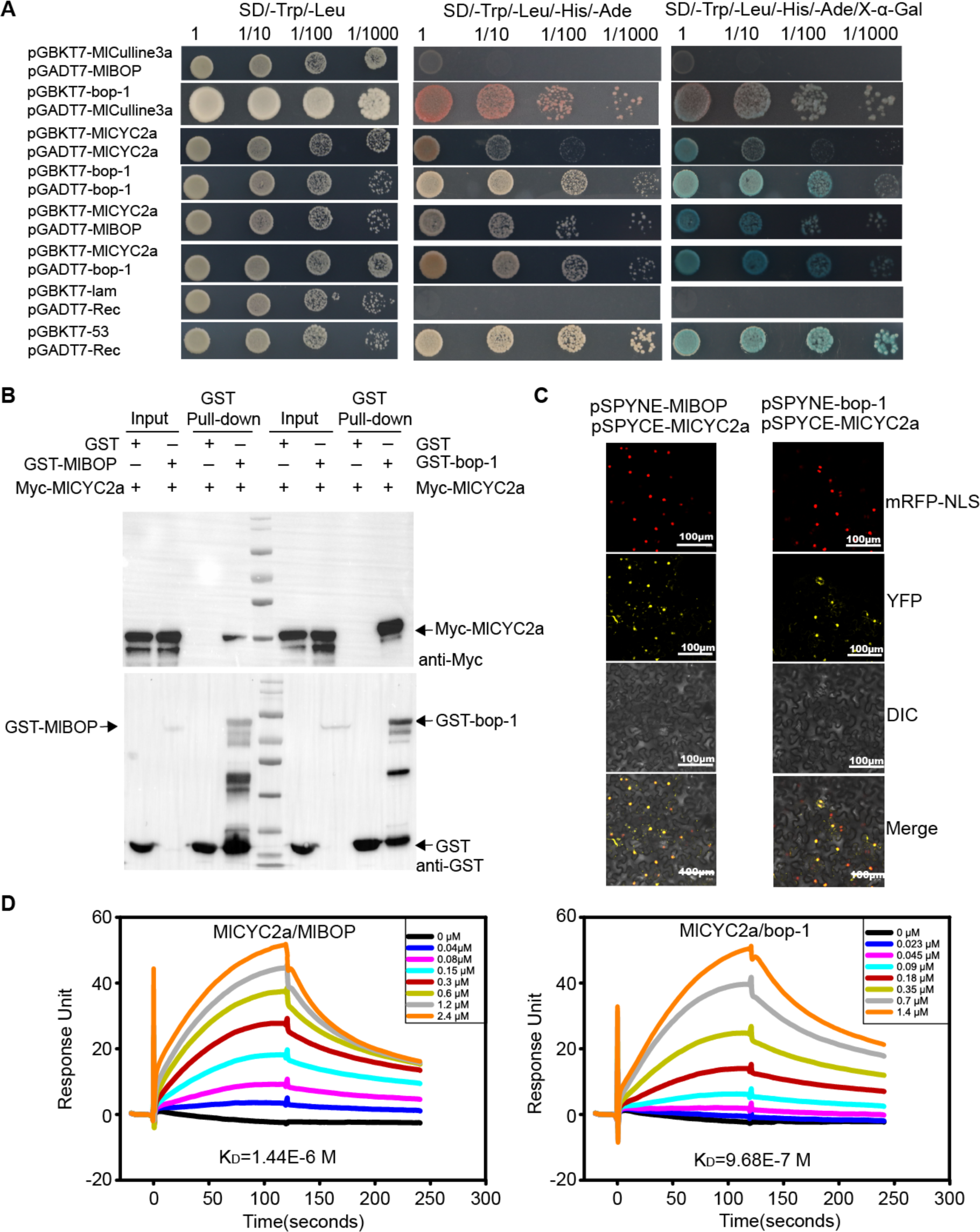
MlBOP interacts with MlCulline3a and MlCYC2a to form a protein complex. (A) Interaction between MlCYC2a or MlCulline3a and MlBOP in yeast cells. SD/- Trp/-Leu indicates Trp and Leu synthetic dropout medium; SD/-Trp/-Leu/-His/-Ade indicates Trp, Leu, His, and Ade synthetic dropout medium. X-*α-*Gal, 5-Bromo-4- chloro-3-indolyl-α-D-galactopyranoside. (B) Interaction between MlBOP or bop-1 and MlCYC2a in an *in vitro* pull-down assay. *In vitro-*translated GST protein was used as the negative control. “Input” indicates protein mixtures before the experiments and “Pull-down” indicates purified protein mixture. “+” indicates presence and “–” indicates absence. (C) Interaction between MlBOP or bop-1 and MlCYC2a in BiFC assay. mRFP-NLS, nuclear marker co-expressing the *35S*:*D53-RFP* construct; YFP, images obtained in the yellow fluorescence channel; DIC, images obtained in bright light; and Merged, overlay plots. (D) SPR sensorgrams demonstrating interaction of MlBOP or bop-1 with immobilized MlCYC2a at different concentrations. Different colors show different concentration of the partner protein, MlBOP or bop-1. The equilibrium dissociation constant (K_D_) value is displayed under each sensorgram, respectively.

Interestingly, we found that the protein-protein interaction between the orthologs of BOP and CYC in the selected species representing different clades of angiosperms is highly conserved: In the two species of Asterids we assayed, we found that AmBOP interacted with both CYC and DICH in snapdragon and CsBOP interacted with some but not all of the CYCs in chrysanthemum, suggesting the functional diversification of CYC after gene duplication in this lineage; In Arabidopsis (Rosids), both AtBOP1 and AtBOP2 interacted with AtTCP1, despite that it bears a radial symmetrical flower; we also found that DaBOP1 and DaBOP2 interacted with DaCYC2b in *Delphinium anthriscifolium*, which belongs to the basal eudicots. Furthermore, we detected the interaction between CeBOP1/2 and CeCYC1/2/3 in *Cymbidium ensifolium* (monocot) (fig. S5, A and C). Taken together, this suggests that BOP-CYC module is highly conserved in monocots and eudicots and the diversification of floral symmetry in different lineages is likely due to the modification of this module or the downstream genes it targets. Intriguingly, BOP from different species interacted with MlCYC2b from *Mimulus* (fig. S5B), despite that some of them did not interact with the corresponding CYC encoded by their own genomes, highlighting the conservative roles of BOP as a pleiotropic regulator of plant development, whereas CYC is less pleiotropic, therefore, more prone to the modification of gene function.

### MlCYC2a functions as a competitor for ubiquitination of MlBOP

To determine whether MlCYC2a can be directly ubiquitinated by the MlCulline3a- MlBOP complex, we conducted *in vitro* ubiquitination assays and reconstituted the MlCulline3a-MlBOP-mediated ubiquitination cascade in *E. coli*. Surprisingly, MlCYC2a was either a poor substrate for ubiquitination or unable to be ubiquitinated by the putative MlCulline3a-MlBOP ligase complex (fig. S6, A and B, output part, anti-MBP). Intriguingly, the presence of MlCYC2a significantly reduced the amount of ubiquitination w/o MlBOP (fig. S6B, lane 5 vs. 1, and lane 7 vs. 6, anti-FLAG), indicating that MlCYC2a may act as a competitor, attenuating the ubiquitination catalyzed by the MlCulline3a-MlBOP ligase complex.

Moreover, the *in vitro* ubiquitination assay showed some ubiquitination occurred in the presence of only E1, E2, E3 and MlCulline3a (fig. S6B, lane 1, anti-FLAG), but the extent of ubiquitination was enhanced in the presence of MlBOP, and reduced when bop-1 was present (fig. S6B, lane 6 vs. 1, and lane 8 vs. 1, anti-FLAG), suggesting that MlBOP can be self-ubiquitinated. Taken together, these results suggest that MlCYC2a is unlikely to be a substrate for the MlCulline3a-MlBOP complex, and the mutation in the BTB domain plays a critical role in MlBOP self- ubiquitination.

### MlBOP and MlCYC2a competitively regulate the self-activation of *MlCYC2a*

Our *in vitro* assays revealed intriguing regulatory relationships between MlBOP and MlCYC2a. Contrary to our initial hypothesis, MlBOP does not seem to ubiquitinate MlCYC2a. Instead, we discovered that MlCYC2a functions as an inhibitor of MlBOP self-ubiquitination, potentially modulating MlBOP homeostasis. This discovery led us to explore alternative regulatory mechanisms between *MlBOP* and *MlCYC2a*.

We first isolated the promoter of *MlCYC2a* (∼ 3kb) in LF10 and identified a putative *cis*-element (P1: GGNCCCNC) matching the consensus CYC*-*like binding site (*33*). Our electrophoretic mobility shift assay (EMSA) and DNA-protein pulldown assay demonstrated that MlCYC2a specifically binds to this P1 element in the *MlCYC2a* promoter *in vitro*, while MlBOP does not (Fig. 5A and Fig. 5B, lane 3&4, output, anti-MBP). The specificity of MlCYC2a binding to the P1 element was further confirmed through a Y1H assay (Fig. 5D) and a chromatin immunoprecipitation (ChIP)-PCR assay using flower buds of stably overexpressing *MlCYC2a* plants (Fig. 5E). These results indicate that MlCYC2a self-activates its own transcription by directly binding to the P1 element in the promoter.

**Fig. 5.**
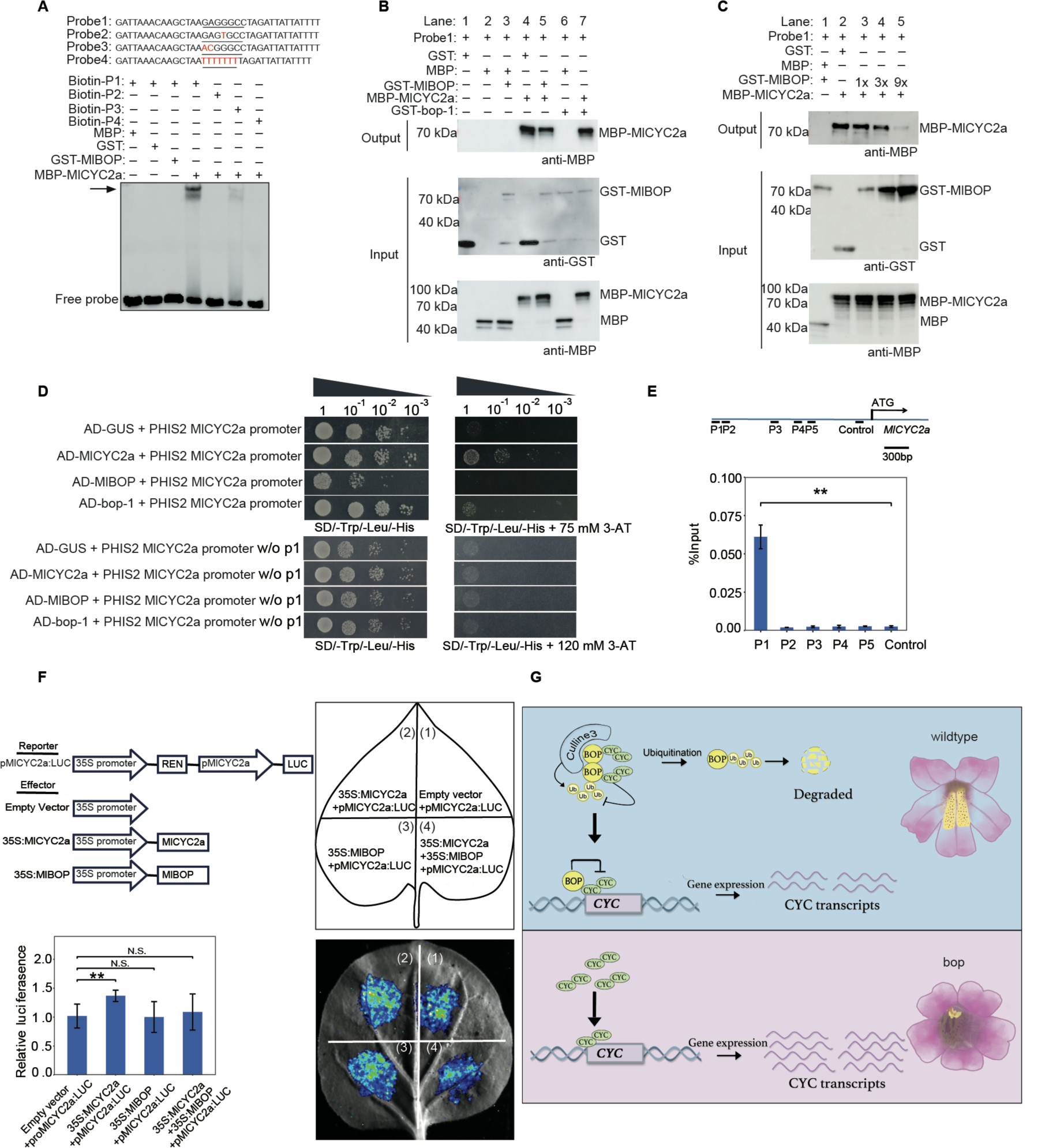
MlBOP interacts with MlCYC2a to modulate the self-activation of *MlCYC2a*. (A) EMSA of interaction between MlCYC2a, MlBOP, and putative MlCYC2a binding sites. The probe was designed from a fragment of the *MlCYC2a* promoter containing the putative MlCYC2a binding motif. Purified GST-tagged MlBOP or MBP-tagged MlCYC2a was incubated with 20 nM wt or mutated biotin-labeled probe. (B) DNA-Protein pulldown assays with the biotinylated P1element from *MlCYC2a* promoter with purified GST-tagged MlBOP or MBP-tagged MlCYC2a. Proteins were incubated for at least 2 hours before pulldown experiments with DNA-bound streptavidin beads. GST and MBP alone were served as negative control and inputs as loading control. (C) MlBOP inhibits the binding of MlCYC2a to the P1 element in a dose-dependent manner. Purified MBP-tagged MlCYC2a was incubated with different amount of GST-tagged MlBOP before pulldown experiments with DNA-bound streptavidin beads. GST and MBP alone were served as negative control and inputs as loading control. (D) Y1H assay showing MlCYC2a specifically binds to the DNA fragment containing the P1 element. Both AD-GUS and AD-bop-1 were used as negative controls. (E) ChIP-PCR assay showing MlCYC2a binding to the *MlCYC2a* promoter in planta. Top, schematic representation of the *MlCYC2a* promoter. Blue line represents the entire length of the *MlCYC2a*; Lines below are the fragments amplified in the ChIP- PCR assay. P1: -2796 to -2676 bp, P2: -2661 to -2560 bp, P3: -1796 to -1660 bp, P4: -1379 to -1275 bp, P5: -1119 to -1015 bp, Control: -263 to -98, relative to the *MlCYC2a* translation initiation codon (ATG). Bottom, ChIP-enrichment of the indicated *MlCYC2a* promoter fragments (P1-P5). P2-P5 were selected due to their fragments containing motifs similar to the putative MlCYC2a binding site and were predicted to bind by other TCP transcription factors in model species. ChIP samples (2 mm flower buds of the *35S*:*MlCYC2a-YFP* transgenic plants) were prepared using the anti-GFP antibody. Error bars represent 1 SD from three biological replicates. (F) Dual-luciferase activity assay showing transactivation of the *MlCYC2a* promoter by MlCYC2a or MlBOP. Top-left, schematic representation of the effector and reporter constructs; Top-right, schematic representation of the tobacco leaves infiltrated with different constructs; Bottom-right, seven independent transfection experiments were performed, and a representative result is shown. Bottom-left, statistical analysis of the LUC/REN value among different constructs infiltrated in the *N. benthamiana* leaves. Error bars represent 1 SD from seven biological replicates. ANOVA test, \*\**p* < 0.01. (G) A working model for the development of flower symmetry by the intricate molecular interaction between *BOP* and *CYC*.

Next, we investigated whether the interaction between MlBOP and MlCYC2a could influence the binding of MlCYC2a to the P1 element. Through a DNA-protein pulldown assay, we found that the presence of MlBOP reduced the amount of MlCYC2a bound to the P1 element (Fig. 5B, Lane 4 vs. Lane 5), indicating that the interaction between MlBOP and MlCYC2a interferes with MlCYC2a binding to its promoter. As increasing amount of MlBOP added to MlCYC2a, less P1 element was detected *in vitro* (Fig. 5C), indicating that MlBOP specifically inhibits the self- activation of *MlCYC2a* in a dose-dependent manner. To further test the repression of MlCYC2a self-activation by MlBOP in planta, we conducted a dual-LUC assay on the same promoter utilized in the Y1H assay, in *N. benthamiana* leaves (Fig. 5F). We found that compared with the control (co-infiltration of empty plasmid with *pMlCYC2a:LUC*), the co-expression of *35S:MlCYC2a* with *pMlCYC2a:LUC* had significantly increased the LUC:REN ratio. By contrast, the co-expression of *35S:MlBOP* with *pMlCYC2a:LUC* did not significantly change the LUC:REN ratio, which is consistent with the lack of a DNA binding domain in *MlBOP*. More importantly, the co-expression of *35S:MlCYC2a* and *35S:MlBOP* with *pMlCYC2a:LUC* had significantly repressed the LUC:REN ratio, indicating that MlBOP can indeed interfere the transcriptional self-activation of *MlCYC2a* in planta.

Taken together, our findings suggest that MlBOP and MlCYC2a competitively regulate the self-activation of MlCYC2a by directly binding to the P1 element in the *MlCYC2a* promoter. The interaction between MlBOP and MlCYC2a appears to modulate the ability of MlCYC2a to activate its own transcription, resulting in a finely tuned regulatory mechanism that influences flower symmetry in *Mimulus*.

## Discussion

Flower symmetry is a crucial evolutionary innovation in angiosperms, and understanding the molecular and developmental mechanisms underlying this trait has been advanced through studies on model organisms like *Antirrhinum majus*. While the spatiotemporal expression pattern of *CYC* has been linked to flower symmetry in various plants (*7, 8, 16, 25, 34–36*), the regulatory mechanisms governing *CYC* expression have remained elusive. In this study, we utilized chemical mutagenesis and functional analysis to identify *BOP* as the causal gene underlying the floral symmetry mutants in *Mimulus*. Our *in situ* hybridization analysis revealed that the spatial and temporal expression of *MlBOP* overlaps with *MlCYC2a*, and probably *MlCYC2b*, suggesting a potential direct regulation of *MlCYC2a* and *MlCYC2b* by *MlBOP*. However, since MlBOP lacks a DNA binding domain, direct binding to the *MlCYC2a* promoter is unlikely. Our *in vitro* assays indicated that MlBOP interacts with both MlCulline3a and MlCYC2a, forming a potential ubiquitination complex to target MlCYC2a for degradation. Surprisingly, MlCYC2a was not ubiquitinated by MlBOP in our *in vitro* ubiquitination assays. Instead, MlBOP was found to be self- ubiquitinated, modulating its own homeostasis. We further demonstrated that *MlCYC2a* self-regulates, and that MlBOP functions as a modulator to fine-tune this MlCYC2a-mediated feedback regulatory loop. This regulatory loop is further complicated by the possible involvement of *MlCYC2b*, which might cross regulates *MlCYC2a*, and vice versa (*33*). This reveals a highly sophisticated and intricate regulatory relationship between *BOP* and *CYC* in *Mimulus* (Fig. 5G). Given the similarity in spatiotemporal patterns of the *MlCYC2a* with those in *Antirrhinum* and other plants (*8, 14, 16, 17, 20*) and the conservative protein-protein interaction between BOP and CYC in many different lineages, this mechanism is likely highly conserved in angiosperms. One particular intriguing observation from our data is that the ability of BOP interacting with CYC seems to be highly conserved, as all of the BOP proteins tested interacted with the MlCYC2b from *Mimulus*, but some of them failed to interact with the cognate CYC proteins encoded by their own genomes. Since BOP regulates an array of plant developmental process, whereas CYC is more specific to the flower development, evolution must have exerted a strong selective pressure on BOP, with less constraint on CYC. Whether this module predates the evolution of the basal angiosperms warrants further investigations by sampling basal angiosperm species. Our findings suggest that evolutionary shifts in flower symmetry across many clades might not necessarily involve *direct* changes of *CYC* alone, but could potentially be influenced by molecular modifications involving BOP and its interacting partners or its downstream targets.

## Acknowledgements

We thank Clinton Morse, Matt Opel, and Meghan Moriarty for extraordinary plant care in the UConn EEB Research Greenhouses. We are grateful to Dr. Yuehua Ma (Central Laboratory of the College of Horticulture, Nanjing Agricultural University for assistance in using the high-resolution confocal laser microscope (ZEISS, LSM800). The plasmids pACYC-RC3, pCDF-Ub, and pETDuet- 1:Flag-WRKY70+GST, were kindly provided by Dr. Xinnian Dong from Duke University. The cDNA of *Delphinium anthriscifolium* was a gift from Dr. Rui Zhang from Northwest A&F University. Dr. Elena Kramer at Harvard University generously provided lab space for B.D. to learn *in situ* hybridization techniques. Wenwen Fan and Yuyang Zhou helped with the drawing art. This project was supported by the Bioinformatics Center of Nanjing Agricultural University.

## Author contributions

B.D., Y.G. and Y-W.Y. planned and designed the research. Y.G., B.D., J.L., J.H., Y.Y., Z.G., Z.Q., Y.Z., Y.K., L.Y., J.W., performed experiments. Q.L. assisted with the SEM. S.C., F.C., provided insightful advice, discussions and support throughout. Y.G., Y-W.Y., B.D. wrote the manuscript with input from all authors.

## Data and materials availability

Genome sequence data have been deposited in the NCBI Sequence Read Archive (SRR26855520). Requests for the materials should be made to B.D.

## Supplementary Materials

### Materials and Methods

#### Plant materials and growth conditions

The *Mimulus lewisii* inbred line LF10 was previously described (*1*). EMS mutants were generated using *M. lewisii* LF10 and *M. verbenaceus* following Owen and Bradshaw (*2*). Plants were grown in the University of Connecticut research greenhouse and Nanjing Agricultural University research greenhouse under natural light supplemented with sodium vapor lamps and LED lights, respectively, ensuring a 16-hr day length.

#### Genomic Analyses for Causal Gene Identification

To identify the causal gene underlying the ML14181, we employed a hybrid strategy that combines the advantages of bulk segregant analysis and genome comparisons between multiple EMS mutants, as described previously (*3*). Briefly, for the ML14181 mutant an F_2_ population was produced by crossing the homozygous mutant (generated in the LF10 background) and the mapping line SL9. DNA samples from 96 F_2_ segregants displaying the mutant phenotype were pooled with equal representation. A small-insert library was then prepared for the pooled sample and was sequenced using an Illumina HiSeq 2000 platform at the University of North Carolina High Throughput Sequencing Facility. These short reads were then mapped to the LF10 genome assembly version 2.0 (http://monkeyflower.uconn.edu/resources/) using CLC Genomics Workbench 7.0 (Qiagen, Valencia, CA). After comparisons to the SNP profiles of previously published mutants, *guideless* (*4*), *rcp1* (*5*), *act1-D* (*6*), and *rcp2* (*7*), we narrowed the causal mutation to a single candidate SNP for ML14181. The allelic ML14132 and NJ01339 mutants were sequenced by Sanger sequencing to detect the mutations.

#### Expression analysis by RT-qPCR

Total RNA was isolated from specified floral tissues using a Quick RNA isolation kit (Huayueyang Biotechnology, Beijing, China) following the manufacturer’s instructions and treated with RNase-free DNase (Vazyme Biotech Co. Ltd., Nanjing, China) to avoid genomic DNA contamination. Reverse transcription was performed to synthesis cDNA from 1μg total RNA using the HiScript II Q Select RT SuperMix (Vazyme Biotech Co. Ltd., Nanjing, China) following the manufacturer’s instructions. Next, cDNA samples were diluted 20-fold before quantitative reverse transcription PCR (RT-qPCR). All RT-qPCRs were performed using the TB Green^®^ Premix Ex Taq™ II (Tli RNaseH Plus) (Takara, Dalian, China) in a CFX96 Touch Real-Time PCR Detection system (Bio-Rad, USA). Samples were amplified for 50 cycles for 95 °C for 15 s and 60 °C for 30 s. *MlUBC* was used as a reference gene to normalize expression levels following the delta-delta Ct method (*1*).

#### Plasmid Construction and Plant Transformation Overexpression plasmids

To generate the *35S*:*MlBOP*, *35S*:*MlCYC2a*, and *35S*:*YFP-MlBOP* constructs, we first amplified the full-length coding sequence (CDS) of *MlBOP* and *MlCYC2a* from the wild-type LF10 cDNA using the Phusion enzyme (NEB, Ipswich, MA), respectively. For each gene, the amplified fragment was cloned into the pENTR/D-TOPO vector (Invitrogen) and then a linear fragment containing the CDS flanked by the attL1 and attL2 sites was amplified using M13 primers. The linear fragments of *MlBOP* and *MlCYC2a* were subsequently recombined into the respective Gateway vector pEarleyGate 100 and pEarleyGate 101 (*8*), which drives transgene expression by the CaMV *35S* promoter. The CDS of *MlBOP* was recombined into the pEarleyGate 100 and 104 vector to complement the *M. lewisii* ML14181 mutant and *M. verbenaceus* NJ01339 mutant phenotypes, respectively.

#### CRISPR-Cas9 plasmid

To knock out the function of *MvBOP*, we undertook a targeted genome editing approach using the Cas9-sgRNA system to recapitulate the *BOP* loss-of-function phenotype in *M. verbenaceus*. Four sgRNA guides were designed to target the first exon of *MvBOP* (sgRNA1:TCAGCGTAGAGGGTCGTCTC; sgRNA2:CAGCTCGGAGCCTCTTCTTC; sgRNA3:GAGCTCCGACCCGTTGAGGA; sgRNA4:CTCGCCGCCGCTAGATCCTT). The 20-bp guide sequences were further verified for their specificity by using a blast algorithm to scan the *M. verbenaceus* v2.0 reference genome (http://mimubase.org/). The sgRNAs were cloned into sgRNAs expression cassettes using overlapping PCR and then ligated by the Golden Gate cloning methods into the binary vector pYLCRISPR/Cas9P35S-B (Addgene#66190) for multiplex gene editing following the previously described protocol (*9*). For genotyping of the transgenic plants, leave tissues of the transgenic plants were collected from each plant and genomic DNA was isolated using the CTAB method. PCR amplification was performed using primers flanking the full length of the *BOP* and Sanger sequencing was used to identify mutations in the *MvBOP* transgenic plants. All plasmids were verified by sequencing before being transformed into *Agrobacterum tumefaciens* strain GV3101 for subsequent stable plant transformation, as previously described (*1*).

#### RNA *in situ* Hybridization

Probe synthesis was performed on cDNA using gene-specific primers (Supp. Table 3) and labelled with DIG RNA Labeling Kit (Cat. No.: 11175025910, Roche Diagnostics GmbH, Mannheim, Germany). Other steps were performed as previously described (*10*). Results were visualized in the Central Laboratory of the College of Horticulture on a Leica DM6 B Upright Microscope using bright field.

#### Yeast Two-Hybrid Assay (Y2H)

Yeast two-hybrid constructs were built using the Matchmaker Gold yeast two-hybrid system (Clontech). The full-length CDSs of *MlCulline3a*, *MlBOP*, *bop-1*, *MlCYC2a*, *MlCYC2b* from *M. lewisii*, AtBOP1, AtBOP2, AtTCP1 from *Arabidopsis thaliana*, *AmBOP*, *AmCYC*, *AmDICH* from *Antirrhinum majus*, *CsBOP*, *CsCYC2a1*, *CsCYC2a2*, *CsCYC2c*, *CsCYC2d*, *CsCYC2e*, *CsCYC2f* from *Chrysanthemum seticuspe*, *DaBOP1*, *DaBOP2*, *DaCYC2a*, *DaCYC2b* from *Delphinium anthriscifolium*, and *CeBOP1*, *CeBOP2*, *CeCYC1*, *CeCYC2*, *CeCYC3* from *Cymbidium ensifolium* were separately recombined into the pGBKT7-BD bait vector, and the pGADT7-AD prey vector using an In-Fusion cloning kit (Clontech). The resultant preys and baits were then co-transformed into *S. cerevisiae* strain Y2H using the lithium acetate method. After inoculating on a selective medium lacking Trp and Leu (-T/-L), the positive colonies were inoculated on a selective medium lacking Trp, Leu, His, and adenine (-L/-T/-H/-A) and grown for 2 days at 28 °C, which were further subjected to X-α-Gal, to identify possible interactions. pGBKT7-53 and pGADT7-Rec were used as positive controls and pGBKT7-lam and pGADT7-Rec were used as negative controls.

#### *In vitro* Pull-down Assay

To test whether MlBOP and bop-1 interact with MlCYC2a, respectively, the CDS of *MlBOP* was cloned by PCR amplification and inserted into the pGEX-5X-1 vector using *BamHI* and *NotI* sites to generate plasmid expressing GST-tagged MlBOP. A point mutation of *MlBOP* (H39L) was generated using the QuickMutationTM site- directed mutagenesis kit (Beyotime, China) to create plasmid expressing GST-tagged MlBOP with the H39L point mutation (bop-1). *MlCYC2a* CDS was cloned by PCR amplification fused with Myc tag (EQKLISEEDL) in N-terminal and inserted into the pET21b vector using *HindIII* site to generate plasmid expressing Myc-tagged MlCYC2a.

Myc-MlCYC2a, GST-MlBOP, GST-bop-1 and GST empty vector were individually expressed in *E.coli* BL21 (DE3) cells. Whole cell lysates (WCL) from *E.coli* BL21 overexpressing MYC-MlCYC2a, GST-MlBOP, GST-bop-1, or GST alone were extracted by ultrasonic with lysis buffer containing 20 mM Tris HCl (pH 7.5), 200 mM NaCl, 1mM EDTA, 1X cocktail of protease inhibitors, 1 mM PMSF (phenylmethylsulfonyl fluoride, Sigma). WCL expressing recombinant GST-tag fusion and MYC-tag fusion proteins were mixed in equal amount and purified with glutathione magnetic agarose beads (Thermo Fisher Scientific, Shanghai, China), washed three times by lysis buffer, separated by 10% SDS-PAGE, transferred onto PVDF membranes (Millipore, USA), and subjected to western blotting following standard protocols. Eluted proteins were probed with anti-MYC (Abmart, China) antibody to identify possible interactions. A similar procedure was followed to test the interaction between AtBOP1 or AtBOP2 and AtTCP1 in an *in vitro* pull-down assay.

#### Bimolecular Fluorescence Complementation (BiFC) Assay

For BiFC assays, the CDSs of *MlBOP* and *bop-1* were cloned into the pSPYNE vector, respectively; while the CDS of the *MlCYC2a* was cloned into the pSPYCE vector (*11*). The resultant vectors were introduced into *Agrobacterium tumefaciens* strain EHA105, and then transiently co-infiltrated in *N. benthamiana* leaves following Ding and Yuan (*12*). The co-expressed *35S:D53-RFP* construct was used as a nuclear marker and visualized by fluorescence microscopy. Fluorescence images were acquired using a high-resolution confocal laser microscope (ZEISS, LSM800) in the Central Laboratory of the College of Horticulture, Nanjing Agricultural University.

#### Surface Plasmon Resonance (SPR) Analysis

SPR experiments were performed using the BIAcore T-200 system at room temperature to quantitatively characterize the binding affinity between the MlCYC2a and MlBOP, MlCYC2a and bop-1. MlCYC2a CDS was cloned by PCR amplification and inserted into the pMBP-C vector using *BamHI* and *NheI* sites to generate plasmid expressing MBP-tagged MlCYC2a. MBP-MlCYC2a was generated from *E.coli* BL21 (DE3) cells and purified using Dextrin Resin 6FF (Sangon Biotech, Shanghai, China) according to the manufacturer’s protocol. GST-MlBOP and GST-bop-1 proteins were purified as mentioned above. MBP-MlCYC2a was immobilized on the CM5 sensor chips (GE Healthcare) via amine coupling. Serially diluted MlBOP/bop-1 proteins in HEPES buffer (0.01 M HEPES, 0.15 M NaCl, 0.5% surfactant P20, 3 mM EDTA, pH 7.4) were followed over the chips. The kinetic data were analyzed with the Biacore T200 Evaluation software using the steady-state affinity model.

#### *E. coli*-based Ubiquitination Assay

To test whether MlCYC2a can be directly ubiquitinated by MlBOP, we performed an *in vitro* ubiquitination assay with a reconstituted *E. coli* system. The ubiquitination reaction was carried out according to the previously described principle of reconstituting basic ubiquitination cascade in *E. coli* (*13*). To reconstitute the complex, CUL3-mediated ubiquitination cascade, the reaction components consisting total of seven proteins (MlCYC2a, MlBOP, CUL3, RBX1, E1, E2 and Ubiquitin) were co-expressed in *E. coli* using a modified Duet vector system (Novagen). Details of the plasmid pACYCDuet-1:RBX1+Myc-CUL3 (pACYC-RC3); pCDFDuet-1:HA-Ub+UBC8+UBA1 (pCDF-Ub) and pETDuet-1:Flag-WRKY70+GST (pET-AdS) were described in a previous study (*14*). Six derivations of pET-AdS plasmid were constructed in this study, including different control plasmids. The CDS of MlCYC2a fused to the MBP tag or MBP tag alone was amplified by PCR and inserted between *NcoI* and *NotI* in MCS-1 of the pETDuet-1: Flag-WRKY70+GST to generate the pETDuet-1: MBP-MlCYC2a+GST and pETDuet-1: MBP+GST. The CDSs of *MlBOP* or *bop-1* fused to GST, were amplified and inserted between *NdeI* and *AvrII* in MCS-II of the pETDuet-1: MBP-MlCYC2a+GST and pETDuet-1: MBP+GST to generate pETDuet-1: MBP-MlCYC2a+GST-MlBOP/bop-1 and pETDuet-1:

MBP+GST-MIBOP/bop-1. The three vectors, pET-AdS, pACYC-RC3 and pCDF-Ub, were co-transformed into the *E. coli* strain BL21(DE3) and ubiquitination reaction was initiated by inducing protein expression with 0.5 mM IPTG for 3 hours at 28 °C. Proteins were extracted by ultrasonic with lysis buffer containing 20 mM Tris HCl (pH 7.5), 200 mM NaCl, 1mM EDTA, 1X cocktail of protease inhibitors, 1 mM PMSF (phenylmethylsulfonyl fluoride, Sigma). Expression of proteins was confirmed with SDS-PAGE on the total lysate. To detect ubiquitination of MBP-MlCYC2a under denaturing conditions, 1% SDS was added to the lysate and heated at 95 °C for 10 min, then diluted 10 times with the lysis buffer and subjected to immunoprecipitation (IP) using a-Dextrin Resin 6FF beads (Sangon Biotech, Shanghai, China). Eluted proteins were subjected to 10% SDS-PAGE and probed with anti-MBP (Transgen, China) and Anti-HA (Invitrogen, USA) antibodies.

#### *In vitro* Ubiquitination Assay

GST-MlBOP, GST-bop-1, MBP-MlCYC2a, MBP and GST proteins were individually expressed in *E.coli* strain BL21 (DE3) and purified with glutathione magnetic agarose beads (for the GST tag, Thermo Fisher Scientific, Shanghai, China) or Dextrin Resin 6FF beads (for the MBP tag, Sangon Biotech, Shanghai, China). The recombinant human ubiquitin-activating enzyme (UBE1), ubiquitin-conjugating enzyme (UbcH5b) and Cullin3/Nedd8/Rbx1 recombinant proteins were purchased from R&D systems (R&D Systems, USA). Ubiquitination reactions were performed as described previously with slight modification (*15*). 100 nM UBE1, 1 mM UbcH5b, 100 nM Cullin3/Nedd/Rbx1, 1uM GST-MlBOP, 1uM MBP-MlCYC2a, were incubated at 30°C for 2 hours in a buffer containing 5 ug flag-ubiquitin, 50 mM Tris-HCl (pH 7.5), 50 mM NaCl, 5 mM MgCl_2_, 5 mM ATP, and 2 mM DTT. The reactions were stopped after 10 min at 95 °C by boiling in DTT-containing SDS loading buffer, and proteins were resolved by 10% SDS-PAGE and immunoblotted.

Anti-Flag antibodies were used for detection of the ubiquitin. Anti-MBP antibodies were used for detection of the MBP proteins. The reactions without E1, E2, or E3, and the reactions with MBP or GST were used as negative controls.

#### Electrophoretic Mobility Shift Assay (EMSA)

GST-MlBOP, GST-bop-1, MBP-MlCYC2a, MBP and GST proteins were purified as mentioned above. Probes of 37-bp in length containing the binding site of the *MlCYC2a* promotor were synthesized and labeled with biotin at their 5’ ends. The probes were also mutated to test the binding specificity of the proteins. Annealed double-stranded probes were incubated with the purified GST-MlBOP and MBP- MlCYC2a in binding buffer (10X Binding Buffer, 1ug/Poly(dI-dC)) for 30 min at 25 °C. DNA-protein complexes were separated by nondenaturing PAGE on ice. EMSA assay was performed using the LightShift™ EMSA Optimization and Control Kit (Thermo Fisher Scientific, Shanghai, China) and Chemiluminescent Nucleic Acid Detection Module Kit (Thermo Fisher Scientific, Shanghai, China), following the manufacturer’s instructions.

#### Yeast One-Hybrid Assay (Y1H)

For Y1H assays, the CDSs of *MlCYC2a*, *MlBOP*, and *bop-1* were inserted into the pGADT7 vector as preys. The promoter fragments of *MlCYC2a* with or without the putative P1 element fragment were cloned into the pHIS2 vector as baits. After cotransformation of the prey and bait into the yeast strain *Saccharomyces cerevisiae* Y187 using the lithium acetate method, the resultant yeast cells were plated onto a selective medium lacking Trp, Leu, and His (SD/-Trp/-Leu/-His). Subsequently, the positive colonies were inoculated on a Trp/-Leu/-His medium supplemented with an appropriate concentration of 3-AT and grown for 3 days at 28°C to identify possible interactions. The CDS of *GUS* (β-glucuronidase) was inserted into the pGADT7 vector as the negative control.

#### DNA-Protein Binding Assay

GST-MlBOP, GST-bop-1, MBP-MlCYC2a, MBP and GST proteins were purified as mentioned above. Biotinylated DNA fragments corresponding to *MlCYC2a* promoter P1 element were generated by polymerase chain reaction. For DNA-protein pulldown, biotinylated P1 fragments were first incubated with streptavidin-bound magnet beads (Thermo Fisher Scientific, Shanghai, China) in binding buffer (50 mM Tris-HCl pH 7.5, 100mM NaCl, 0.05% Nonidet P40 and 1mM EDTA) for 2 h at 4 °C, then washed three times in binding buffer. Proteins were added to DNA-bound beads and the mixture was rotated in a 4 °C cold room for 2 h. Beads were washed three times with binding buffer; proteins were stripped off the beads by boiling with 2 x SDS buffer and then subjected to SDS-PAGE. The gel was transferred onto PVDF membranes (Millipore, USA), and subjected to western blotting following standard protocols.

Eluted proteins were probed with anti-MBP or anti-GST (Abmart, China) antibody to identify possible interactions.

#### ChIP-PCR Assay

For ChIP assays, 1.5g of ∼2 mm flower buds of *35S*:*MlCYC2a-YFP* were cross-linked in polyformaldehyde. The chromatin was sheared to an average of 500 bp by sonication, and immunoprecipitated with GFP recombinant rabbit monoclonal antibody (Thermo Fisher Scientific). Subsequently, the enriched DNA fragments were examined by RT-qPCR.

#### Phylogenetic Analysis

Multiple sequence alignment of BOP and related proteins was performed using AliView (*16*). To identify putative orthologous of *CYC* in *Mimulus*, *CYC*, *DICH* from snapdragon and multiple related proteins from other species were aligned to perform phylogenetic analysis. Maximum likelihood analysis was conducted in MEGA (*17*) with the default setting, except for a bootstrap value of 10,000.

#### Dual-LUC Reporter Assay

For Dual-LUC reporter assay, the promoter fragment of *MlCYC2a* was cloned into the pGreenII 0800-LUC vector (*18*) to generate the reporter construct. The CDSs of the *MlBOP* and *MlCYC2a* were cloned into the *pORE-R4* vector (*19*) under the control of the *35S* promoter to generate the effector constructs. Subsequently, the *Agrobacterium tumefaciens* containing the reporter constructs and effector constructs, respectively, were transiently co-infiltrated in *N. benthamiana* leaves. The LUC-to-REN activity ratio was measured using the Infinite M200 luminometer (Tecan, Mannerdorf, Switzerland) with the Dual-Glo^®^ Luciferase Assay System (Promega, Beijing, China). All primers are listed in Supplement Table 3.

#### Scanning Electron Microscopy

Flower buds were fixed overnight in Formalin-Acetic-Alcohol (FAA) at 4°C, dehydrated for 30 min through a 50%, 60%, 70%, 95%, and 100% alcohol series. Samples were then critical-point dried, mounted, and sputter coated before being observed using a NOVA NanoSEM with Oxford EDX at 35 kV at UConn’s Bioscience Electron Microscopy Laboratory.

---

**Figure S1.**
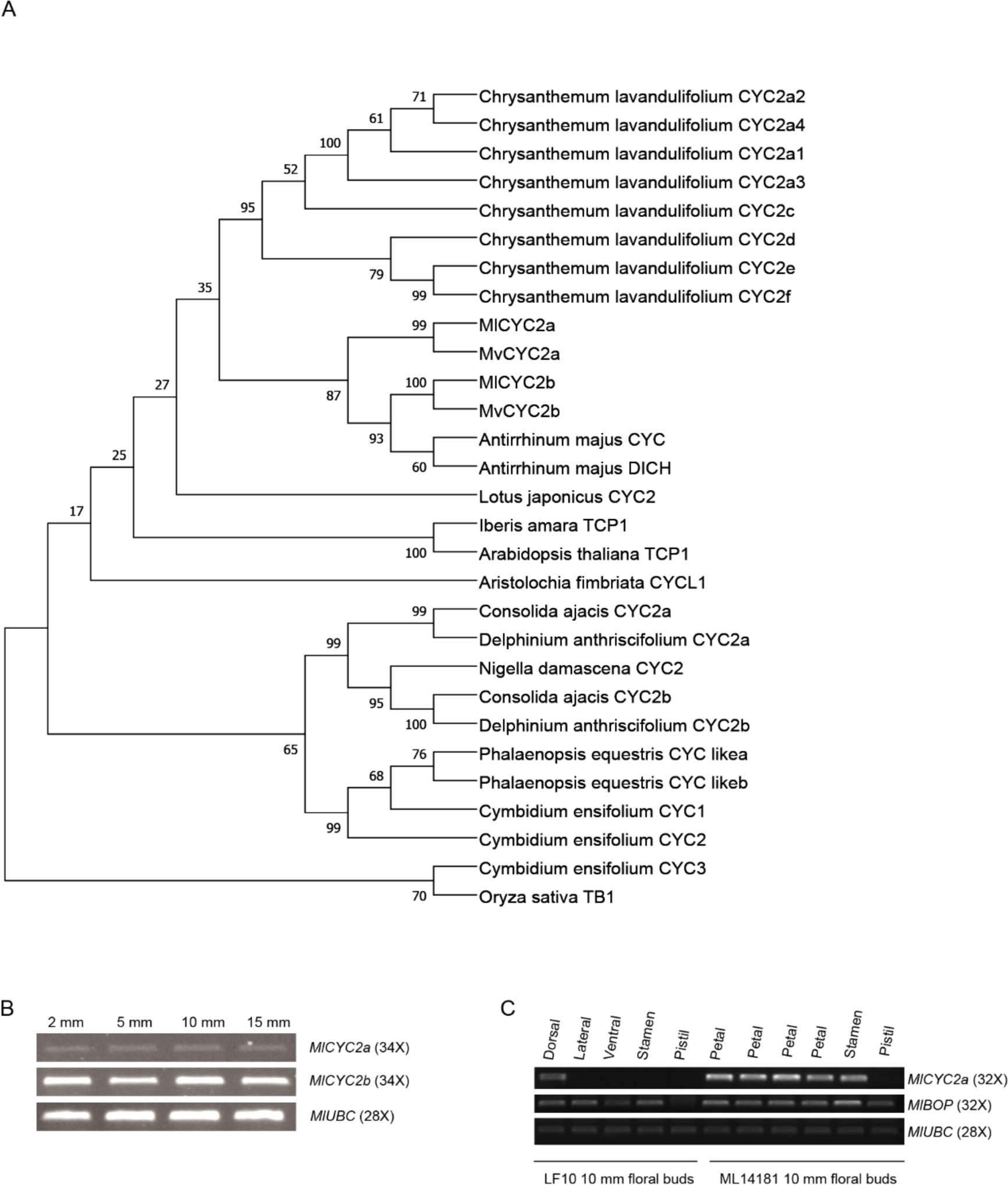
**(A)** Phylogenetic analysis of *CYC.* All *Mimulus* sequences were retrieved from the *Mimulus* genome database (http://mimubase.org/) under the following accession numbers: MlCYC2a, ML4G049700.1; MlCYC2b, ML3G247900.1; MvCYC2a, MV4G047200.2; MvCYC2b, MV3G243500.1. The rest of the sequences were retrieved from GenBank or UniProtKB/Swiss-Port (*Oryza sativa* TB1, BAC99350.1; *Antirrhinum majus* CYC, O49250.1; *Antirrhinum majus* DICH, Q9SNW8.1; *Lotus japonicus* CYC2, ABB36472.1; *Iberis amara* TCP1, ABV57375.1; *Arabidopsis thaliana* TCP1, NP_001077781.1; *Phalaenopsis equestris* CYC likea, XP_020584437.1; *Phalaenopsis equestris* CYC likeb, XP_020584438.1; *Aristolochia fimbriata* CYCL1, AJE30095.1; *Consolida ajacis* CYC2a, UPY89970.1; *Consolida ajacis* CYC2b, UPY89971.1; *Delphinium anthriscifolium* CYC2a, UKB40098.1; *Delphinium anthriscifolium* CYC2b, UKB40099.1); *Cymbidium ensifolium* CYC1, OR823955; *Cymbidium ensifolium* CYC2, OR823956; *Cymbidium ensifolium* CYC3, OR823957. The chrysanthemum sequences were retrieved from the Chrysanthemum Genome Database with the following accession numbers: *Chrysanthemum lavandulifolium* CYC2a1; EVM0031596.1; *Chrysanthemum lavandulifolium* CYC2a2, EVM0062019.1; *Chrysanthemum lavandulifolium* CYC2a3, EVM0012480.1; *Chrysanthemum lavandulifolium* CYC2a4, EVM0076812.1; *Chrysanthemum lavandulifolium* CYC2c, EVM0019255.1; *Chrysanthemum lavandulifolium* CYC2d, EVM0064455.1; *Chrysanthemum lavandulifolium* CYC2e, EVM0029187.1; *Chrysanthemum lavandulifolium* CYC2f, EVM0013168.1. Bootstrap values are shown along the branches. The tree was rooted by midpoint rooting. **(B)** *MlCYC2a* and *MlCYC2b* expression patterns in flower buds at different developmental stages revealed by semi-quantitative PCR. **(C)** *MlCYC2a* and *MlBOP* expression patterns in different floral organs of the dissected wild type and ML14181 mutant floral buds at 10-mm stage revealed by semi-quantitative PCR.

**Figure S2.**
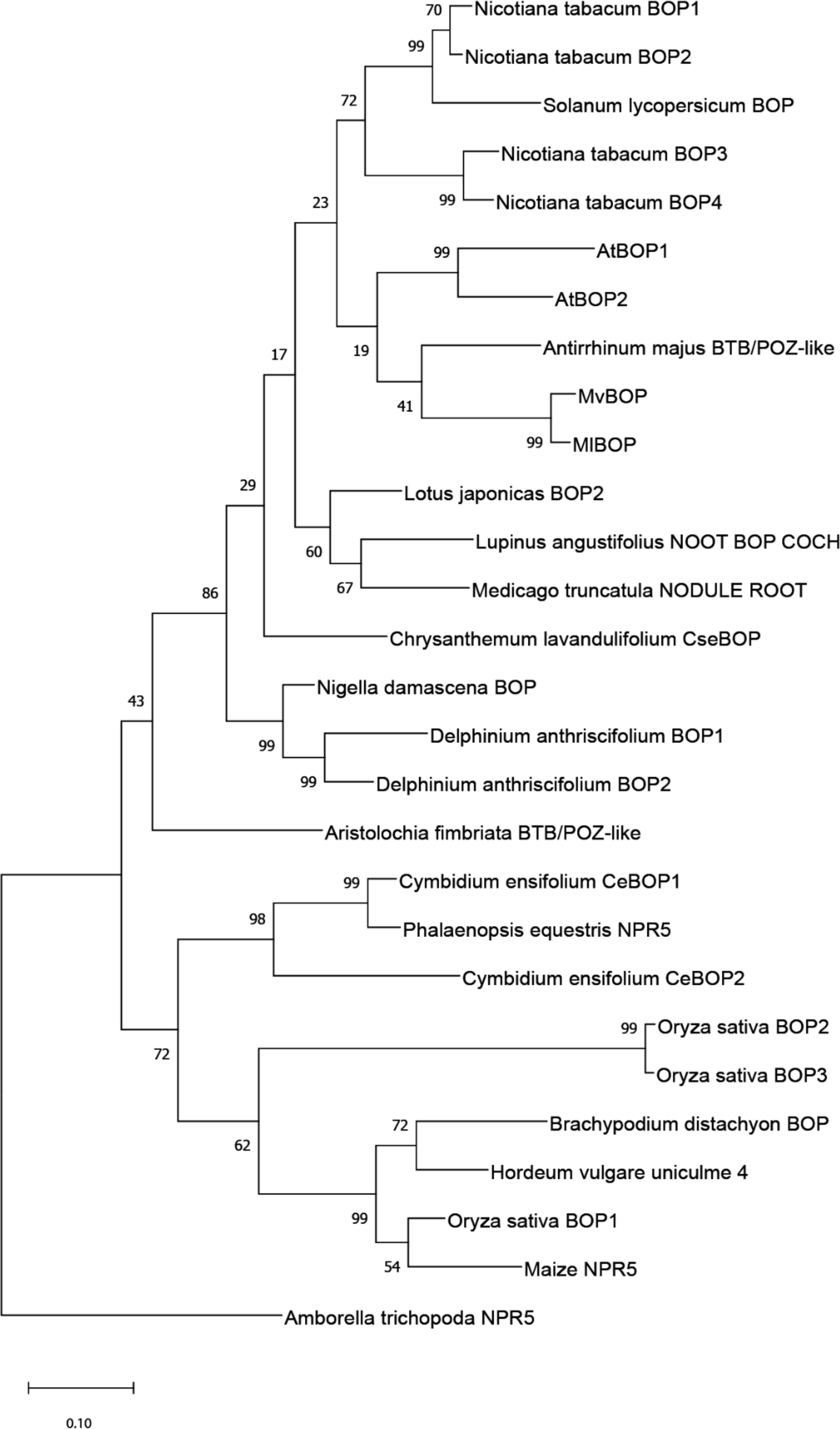
Maximum Likelihood (ML) phylogenic tree of BOP and related proteins. All *Mimulus* sequences were retrieved from the *Mimulus* genome database (http://mimubase.org/) under the following accession numbers: MlBOP, ML8G244400.1; MvBOP, MV8G351700.1. The rest of the sequences were retrieved from GenBank or UniProtKB/Swiss-Port (NtBOP1, XP_016498803.1; NtBOP2, XP_016515209.1; NtBOP3, NP_001312808.1; NtBOP4, NP_001312564.1; *Solanum lycopersicum* BOP, XP_004237330.1; AtBOP1, AT3G57130; AtBOP2, AT2G41370; *Lotus japonicas* BOP2, XP_057437963.1; *Lupinus angustifolius* NOOT BOP COCH, XP_019458812.1; *Medicago truncatula* NODULE ROOT, AET34786.1; *Antirrhinum majus* BTB/POZ-like, Am03g35310 from the Snapdragon Genome Database; *Chrysanthemum lavandulifolium* ClBOP, OR826817; *Nigella damascena* BOP, OR823960; *Delphinium anthriscifolium* BOP1, OR823961; *Delphinium anthriscifolium* BOP2, OR823962; *Aristolochia fimbriata* BTB/POZ-like, KAG9443643.1; *Phalaenopsis equestris* NPR5, XP_020575938.1; *Oryza sativa* BOP1, NP_001388418.1; *Oryza sativa* BOP2, NP_001410083.1; *Oryza sativa* BOP3, NP_001410243.1; *Brachypodium distachyon* BOP, XP_003565065.1; *Cymbidium ensifolium* CeBOP1, OR823958; *Cymbidium ensifolium* CeBOP2, OR823959; *Hordeum vulgare* uniculme 4, AHY22620.1; Maize NPR5, NP_001140651.1; *Amborella trichopoda* NPR5, XP_006843393.1). Bootstrap values are shown along the branches. The tree was rooted by midpoint rooting.

**Figure S3.**
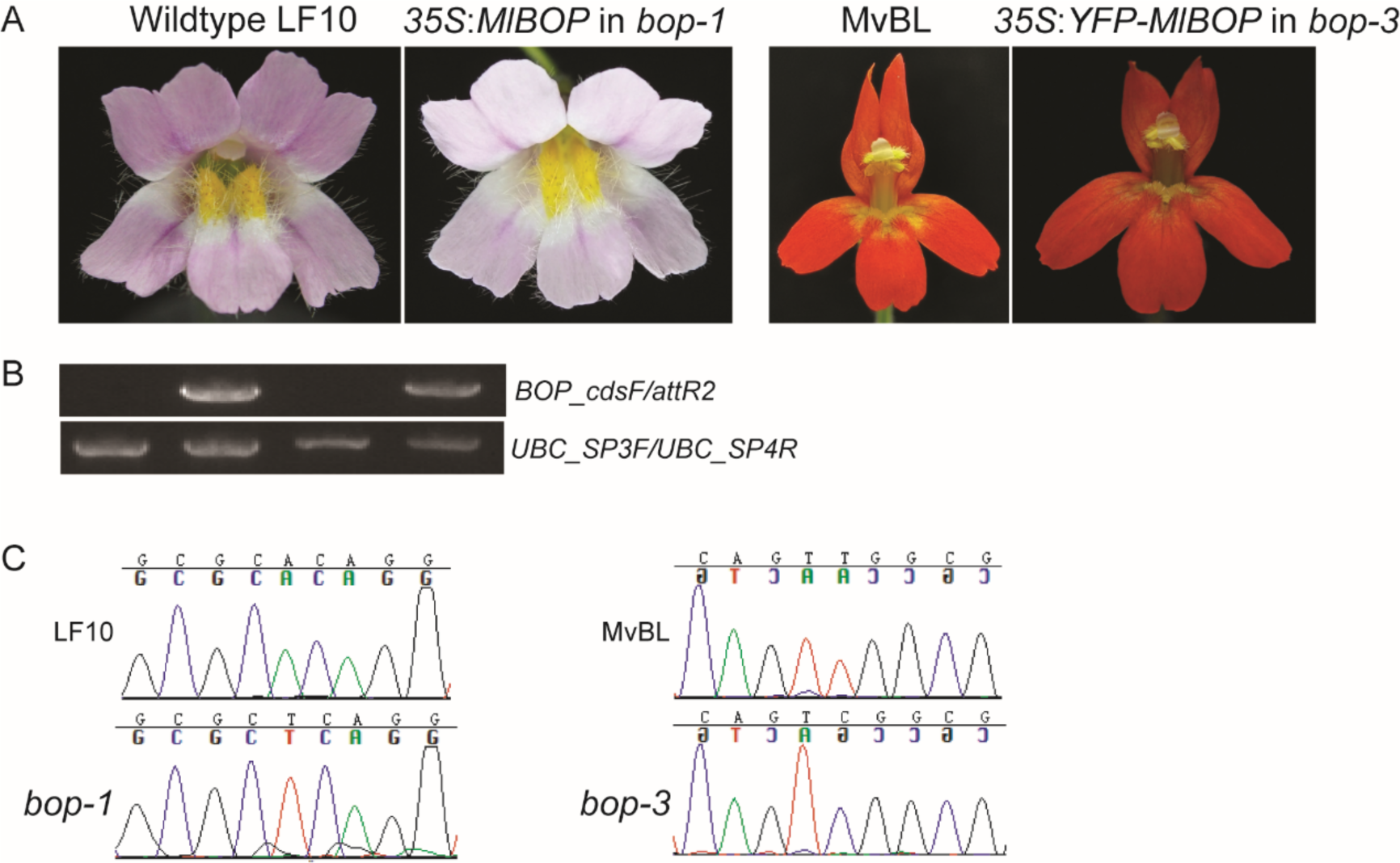
(A) Overexpression of *MlBOP* and *YFP-MlBOP* complements the *bop-1* and *bop-3* mutants, respectively. (B) RT-PCR to validate the complemented F_2_ segregants contains the transgene in them. The top gel examines the presence or absence of the transgene in the samples. The bottom gel is a positive control of PCR using a housekeeping gene UBC. The order of the samples in the gels from left to right follows the same order as that in (A). (C) Sanger sequencing confirms that the same complemented F_2_ segregants are indeed in the mutant backgrounds.

**Figure S4.**
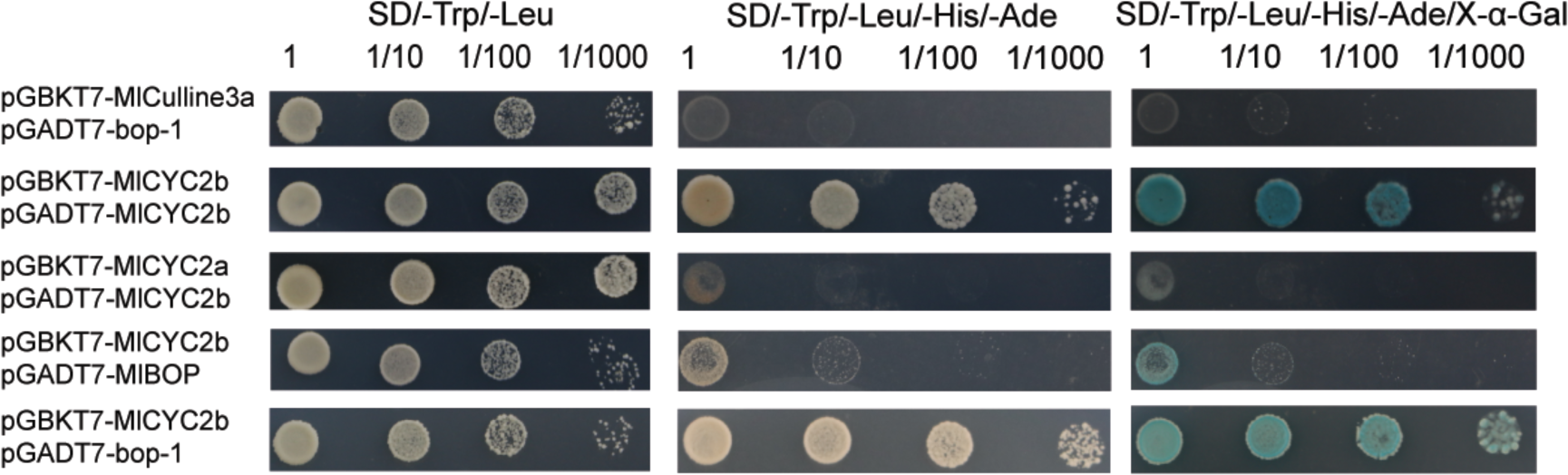
Protein-protein interaction between MlCulline3a, MlBOP, bop-1, MlCYC2a, and MlCYC2b as revealed by Y2H. SD/-Trp/-Leu indicates Trp and Leu synthetic dropout medium; SD/-Trp/-Leu/-His/-Ade indicates Trp, Leu, His, and Ade synthetic dropout medium. X-*α-*Gal, 5-Bromo-4-chloro-3-indolyl-α-D-galactopyranoside.

**Figure S5.**
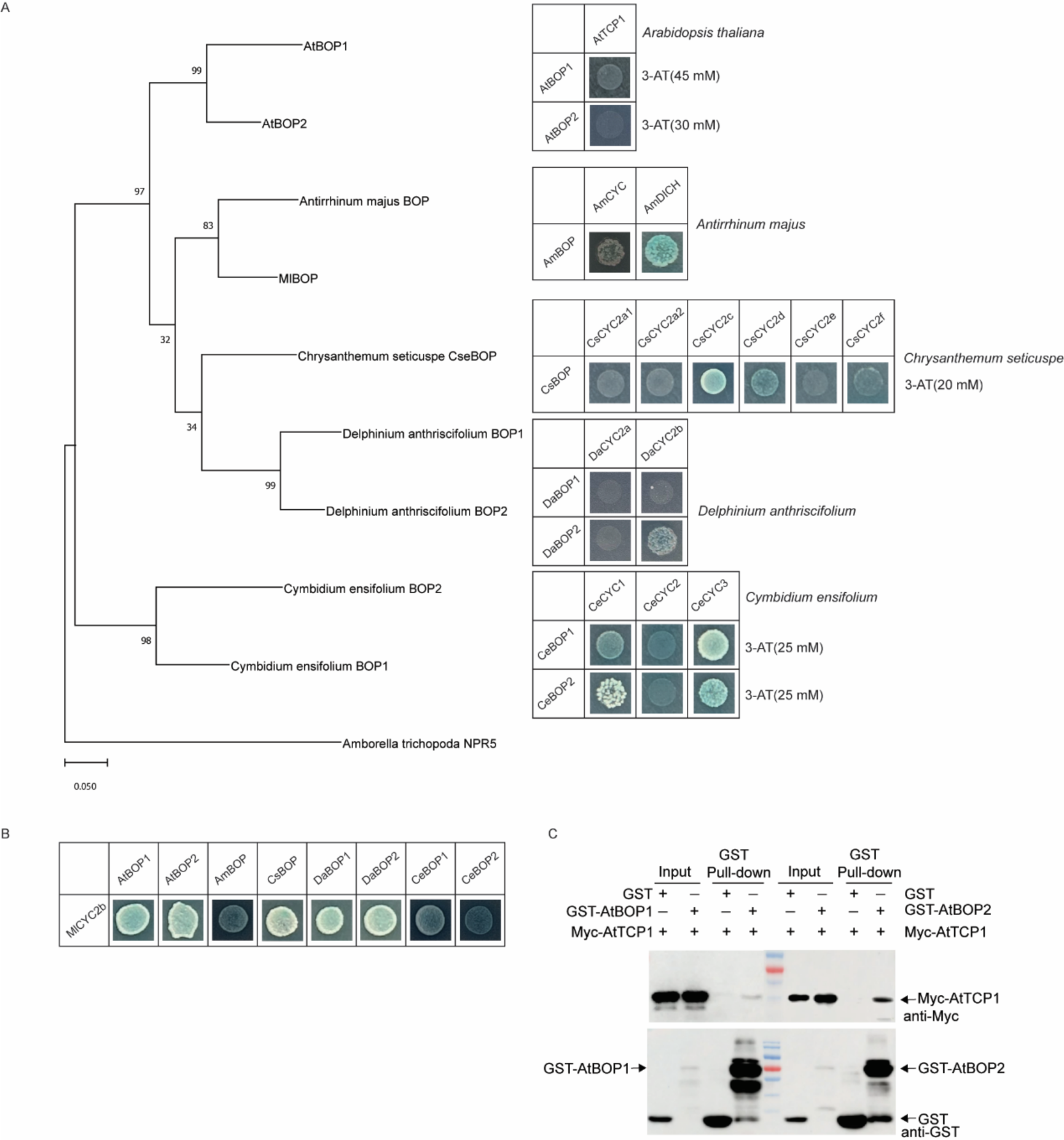
(A) Y2H analysis of the BOP and the cognate CYC from species belonging to different phylogenetic clades. Different concentrations of 3-AT were used to suppress the yeast self-activation. (B) BOP of the selected species from different phylogenetic clades all interacted with MlCYC2b. SD/-Trp/-Leu indicates Trp and Leu synthetic dropout medium; SD/-Trp/-Leu/-His/-Ade indicates Trp, Leu, His, and Ade synthetic dropout medium. X-*α-*Gal, 5-Bromo-4-chloro-3-indolyl-α-D-galactopyranoside. (C) Interaction between AtBOP1 or AtBOP2 and AtTCP1 in an *in vitro* pull-down assay. *In vitro-*translated GST protein was used as the negative control. “Input” indicates protein mixtures before the experiments and “Pull-down” indicates purified protein mixture. “+” indicates presence and “–” indicates absence.

**Figure S6.**
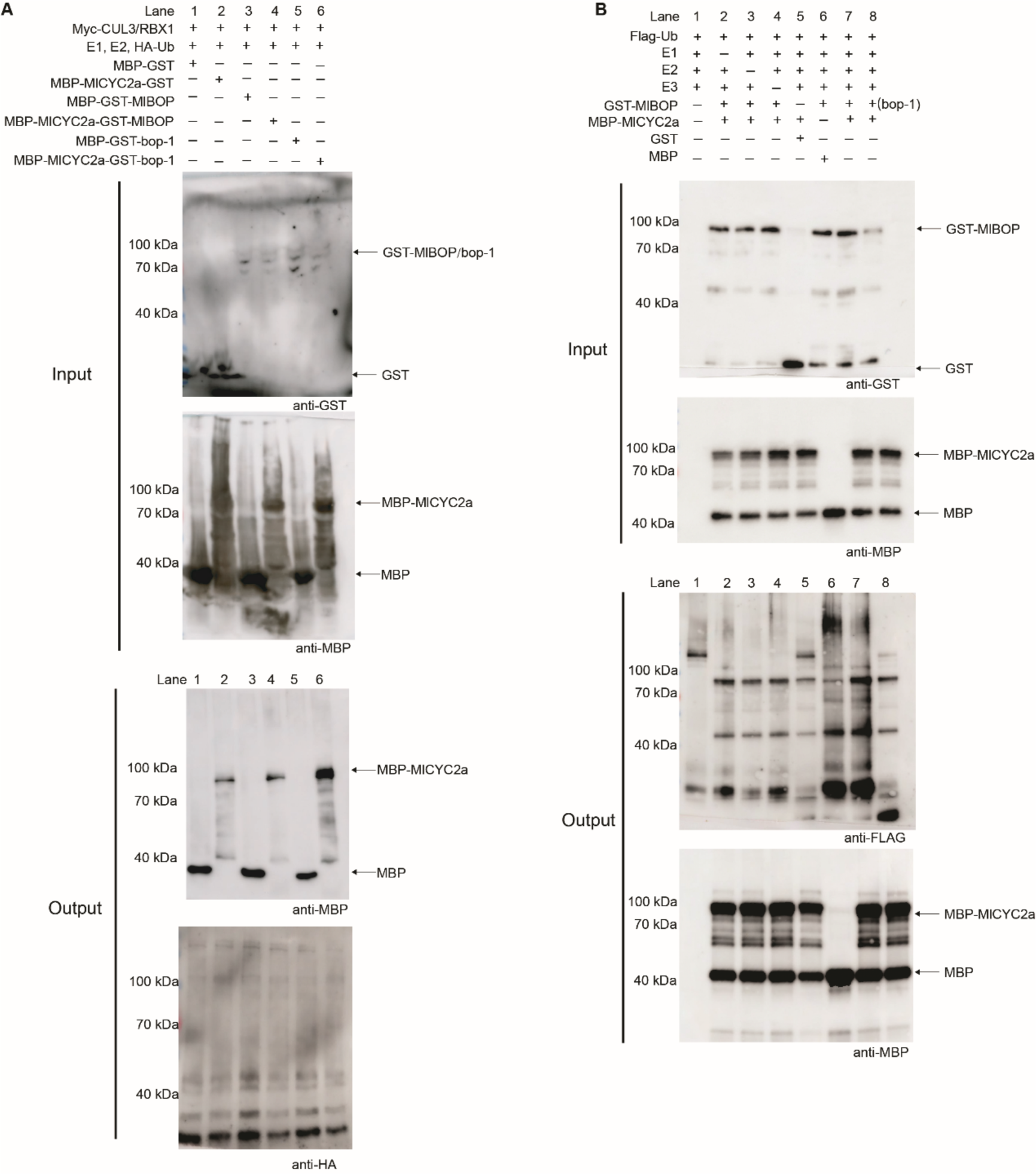
*In vitro* ubiquitination assay of MlBOP and MlCYC2a. (A) Ubiquitination assay of MlCYC2a by the MlBOP-MlCulline3a ubiquitin ligase reconstituted in *E. coli*. MBP-MlCYC2a was immunoprecipitated under denaturing conditions. (B) *In vitro* ubiquitination assay of the recombinant GST-MlBOP and MBP- MlCYC2a in the presence or absence of E1 (UBE1), E2 (UbcH5b), Cullin3 E3 ligase complex (Human CUL3/NEDD8/RBX1) and Flag-ubiquitin (the + indicates added sample). Ubiquitination was detected by immunoblotting with an anti-Flag antibody. MBP-MlCYC2a were detected by immunoblotting with anti-MBP antibody. Inputs are shown using anti-GST and anti-MBP antibodies. MBP and GST were used as negative controls.

**Table S1.** Comparisons of floral traits between the wild type and the mutants (mean±1SD)

|  | No. of<br>calyx | No. of extra tissues<br>grown between<br>calyx and corolla | No. of<br>petals | No. of<br>stamens | No. of extra<br>tissues grown<br>between petals<br>and stamens | 1025<br>1026<br>No. of<br>pistil<br>lobes |
| --- | --- | --- | --- | --- | --- | --- |
| wildtype | 5 | 0 | 5 | 4 | 0 | 2 |
| ML14181<br>(n=20) | 4 | 2.7±0.98 | 4 | 3.7±0.88 | 0.6±0.83 | 2.8±0.85 |
| ML14132<br>(n=20) | 5 | 3.6±1.05 | 5 | 3.4±0.81 | 2.6±0.76 | 2 |

**Supplement Table 2.** Candidate SNPs from the mutant genome comparisons. The SNP highlighted in bold is the causal mutation.

| LF10g_v2.0 | Position | Mutant | Wild-type | Annotation |
| --- | --- | --- | --- | --- |
| LF10_chr1 | 9464985 | T | C | Protein-coding |
| LF10_chr2 | 15356616 | TAATTC<br>TTCAAA | T | Protein-coding |
| LF10_chr2 | 49941784 | T | A | Protein-coding |
| LF10_chr3 | 23018872 | C | G | Protein-coding |
| LF10_chr4 | 13116461 | C | T | Non-coding intergenic sequence |
| LF10_chr4 | 25610766 | G | A | Protein-coding |
| LF10_chr6 | 5331114 | T | C | Protein-coding |
| LF10_chr6 | 7402455 | G | T | Non-coding intergenic sequence |
| LF10_chr6 | 16112976 | A | G | Non-coding intergenic sequence |
| LF10_chr6 | 31264122 | A | G | Protein-coding |
| LF10_chr6 | 37788399 | A | G | Protein-coding |
| LF10_chr6 | 40772786 | T | C | Non-coding intergenic sequence |
| LF10_chr6 | 43034822 | A | T | Protein-coding |
| LF10_chr6 | 48741069 | A | T | Non-coding intergenic sequence |
| LF10_chr6 | 55456975 | G | A | Protein-coding |
| LF10_chr6 | 56252271 | G | A | Non-coding intergenic sequence |
| LF10_chr6 | 58847009 | C | T | Protein-coding |
| LF10_chr6 | 62506135 | A | AG | Protein-coding |
| LF10_chr6 | 63755755 | C | A | Protein-coding |
| LF10_chr6 | 66625657 | G | T | Protein-coding |
| LF10_chr6 | 67722256 | G | T | Protein-coding |
| LF10_chr6 | 67722293 | CT | C | Protein-coding |
| **LF10_chr8** | **37127265** | **T** | **A** | **Protein-coding** |
| LF10_chr8 | 38693100 | C | T | Protein-coding |

**Supplement Table 3.**
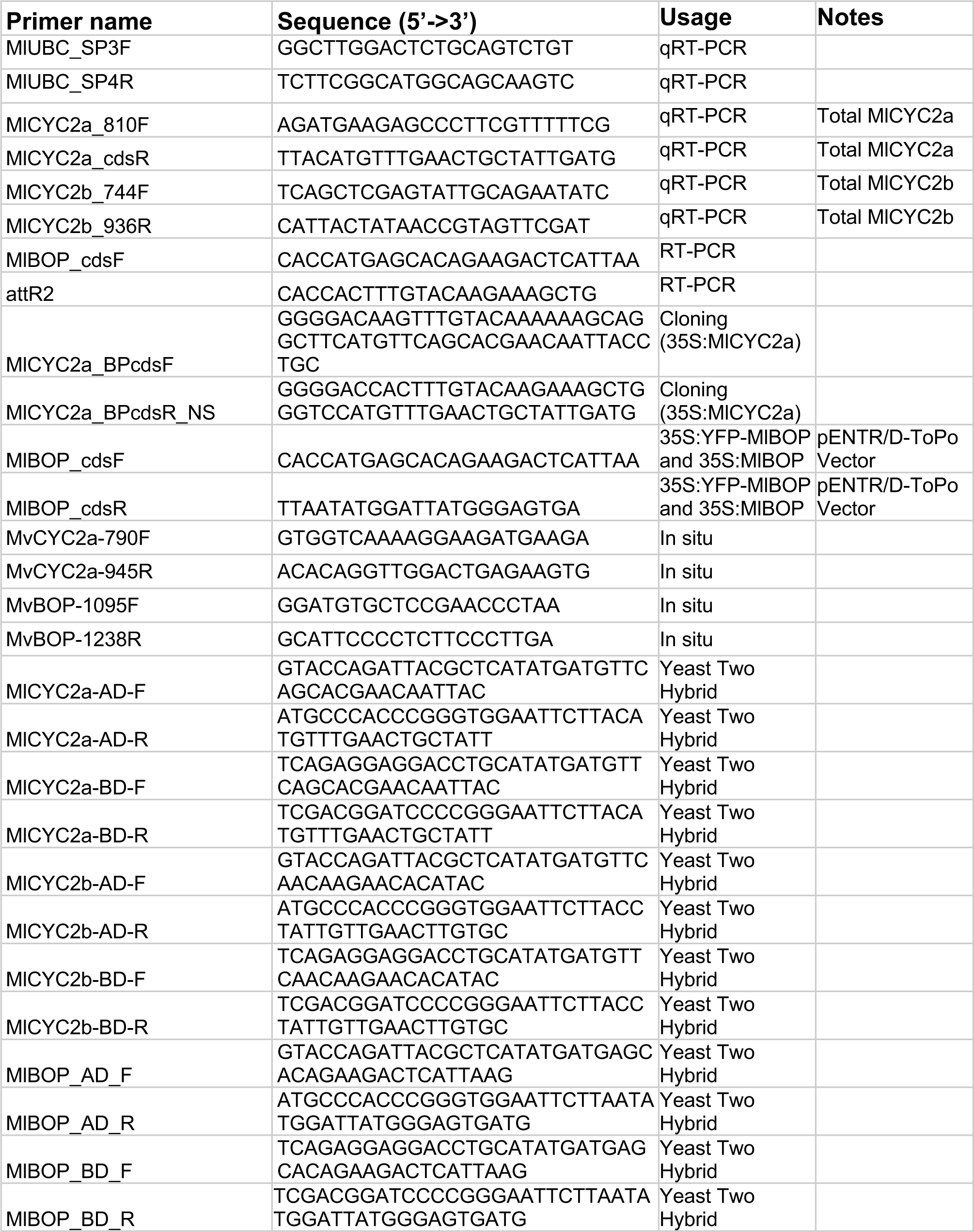
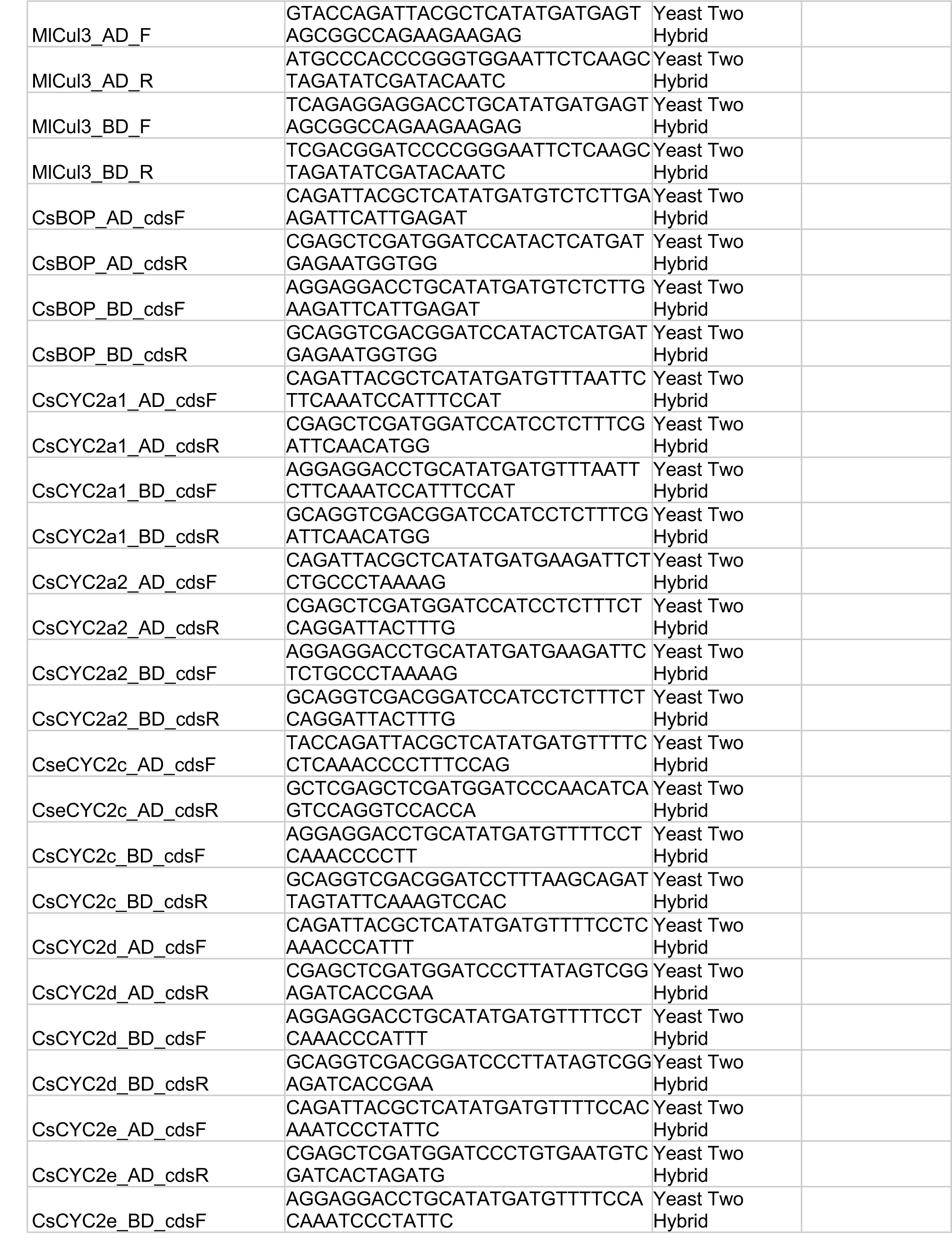
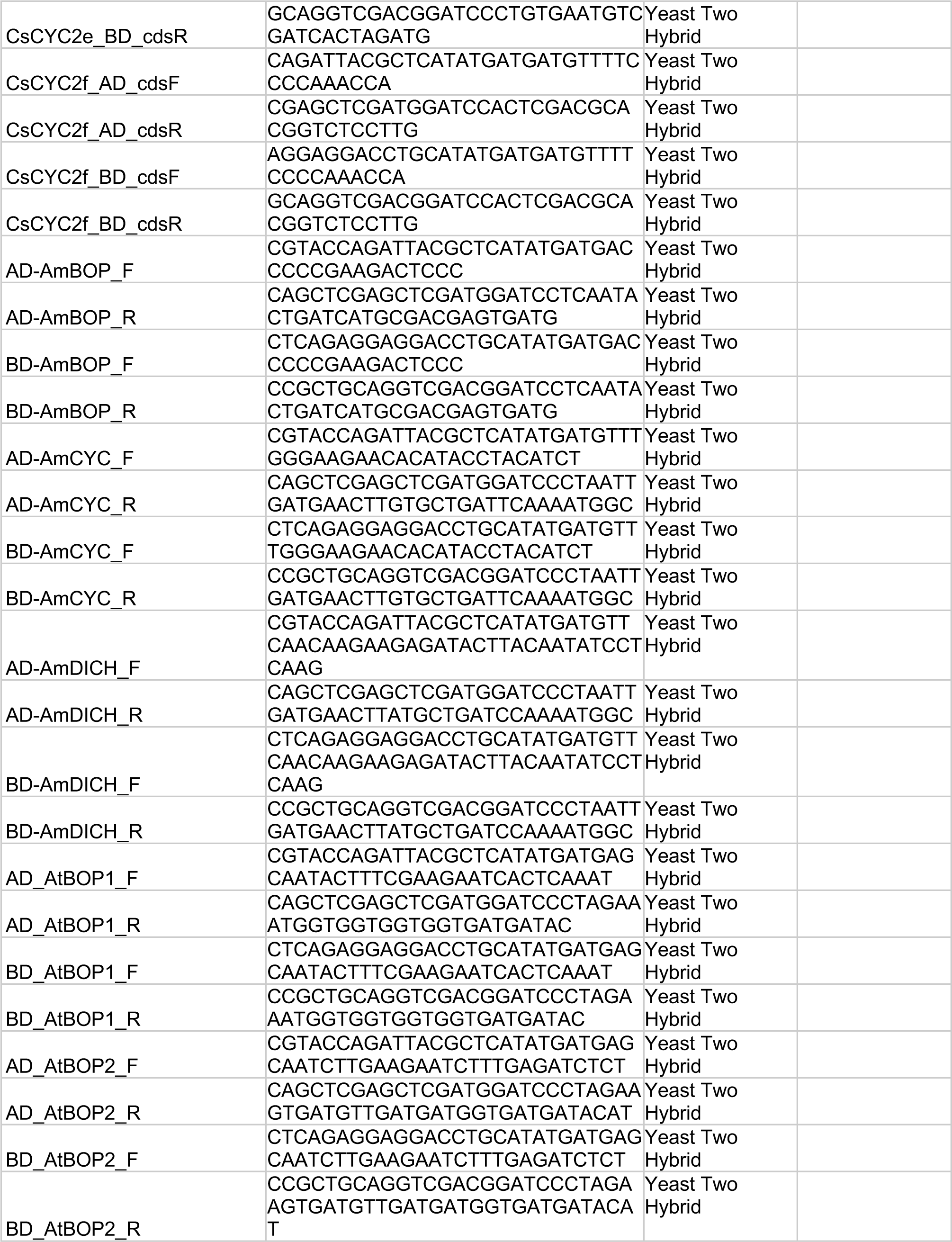
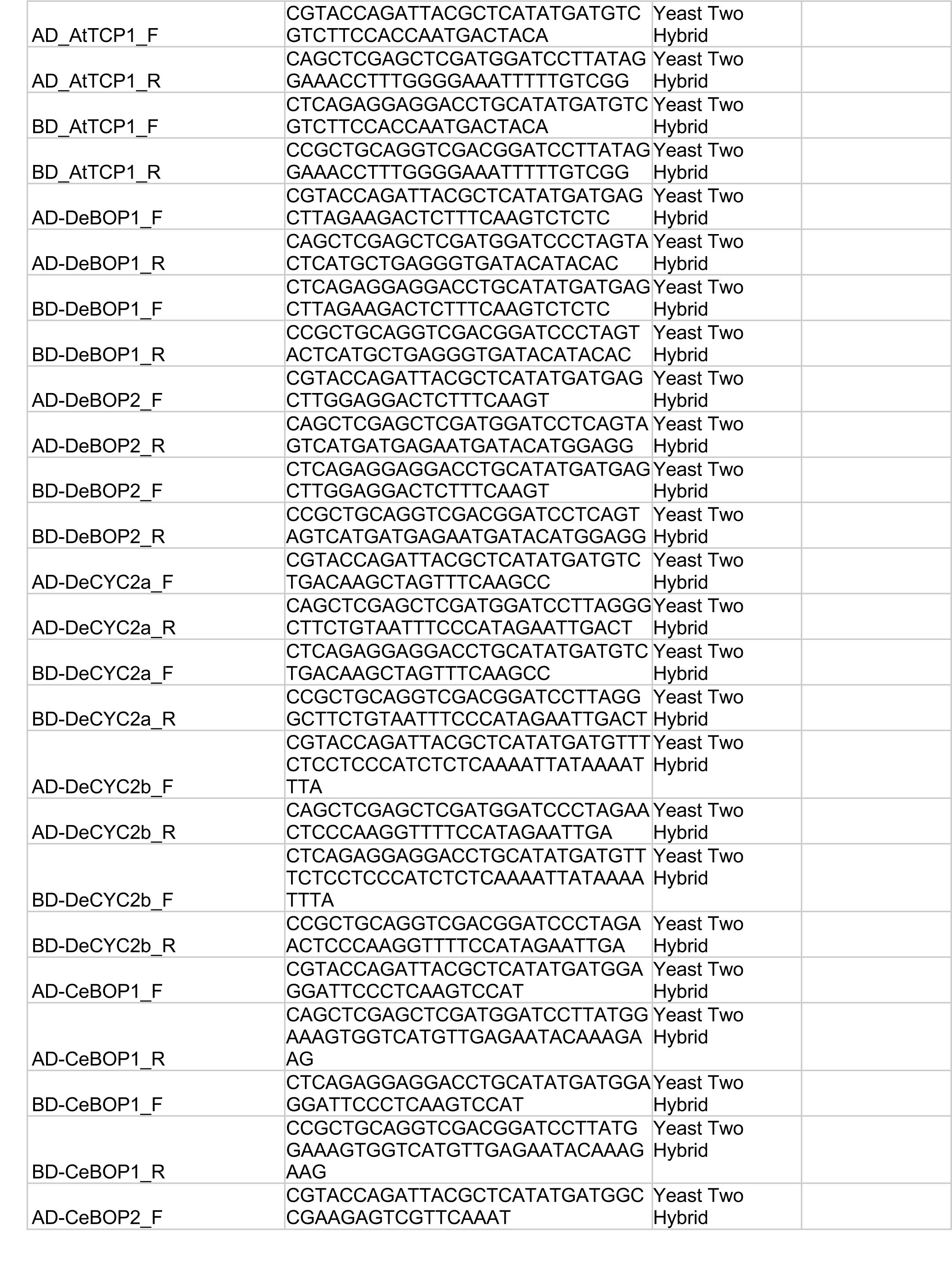
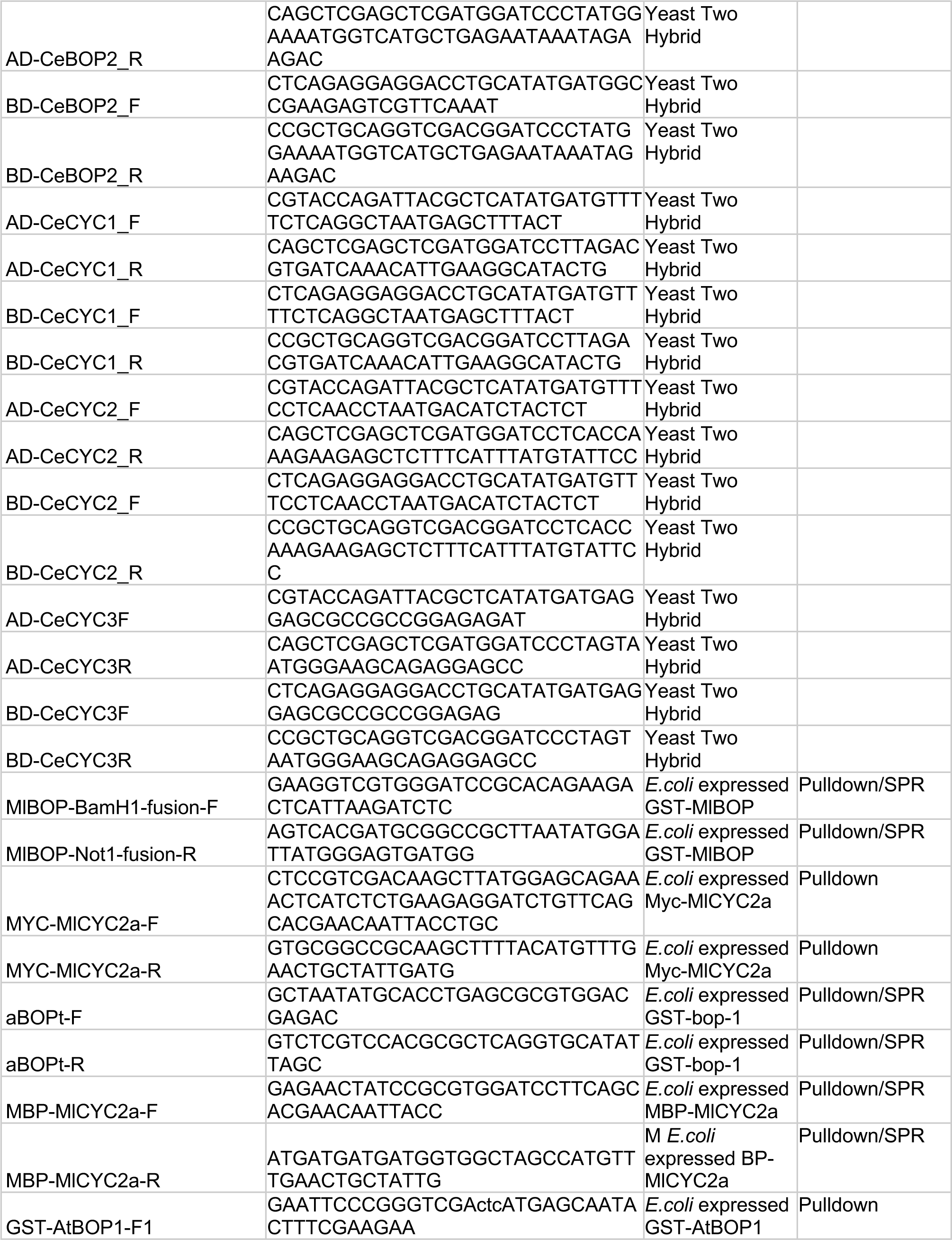
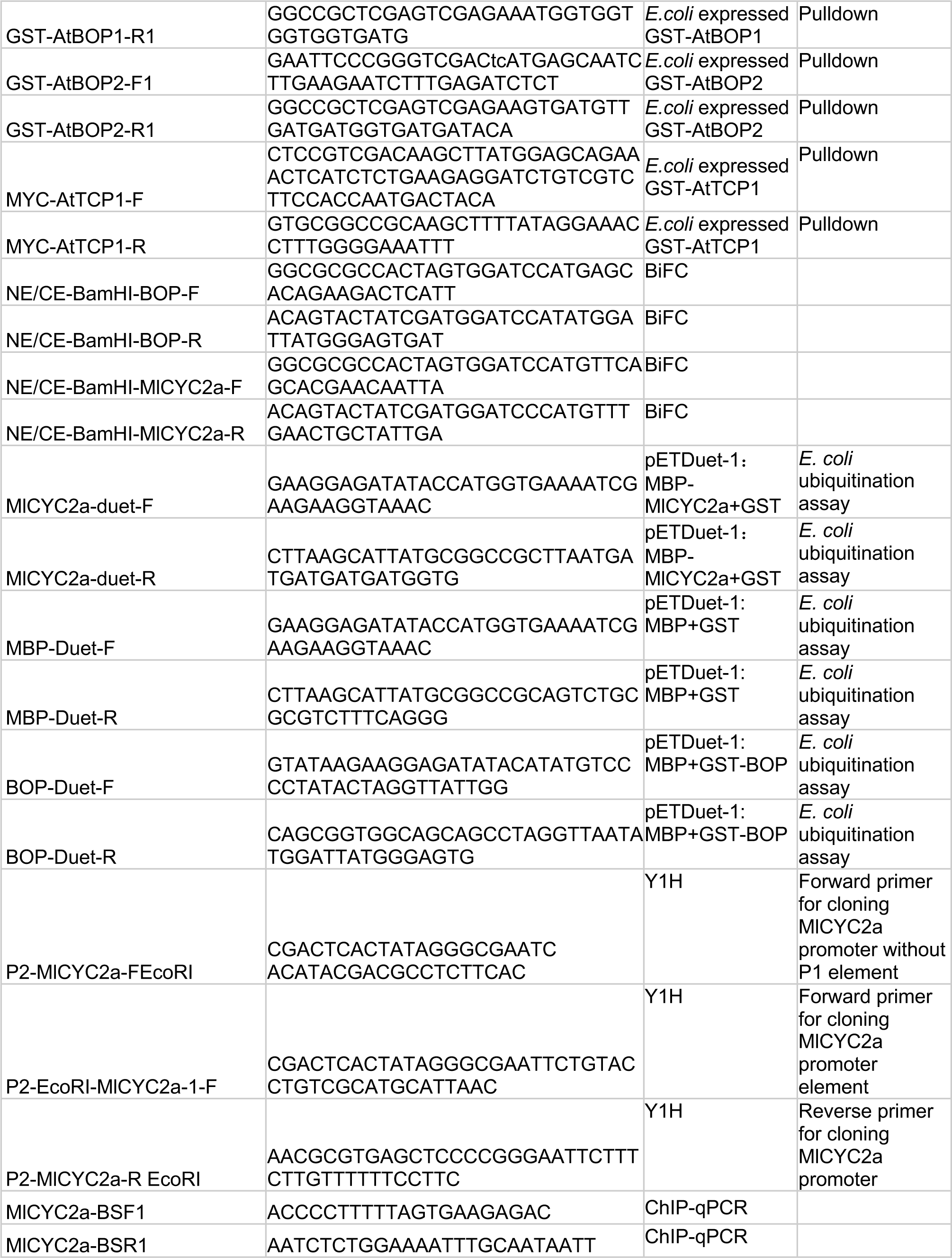
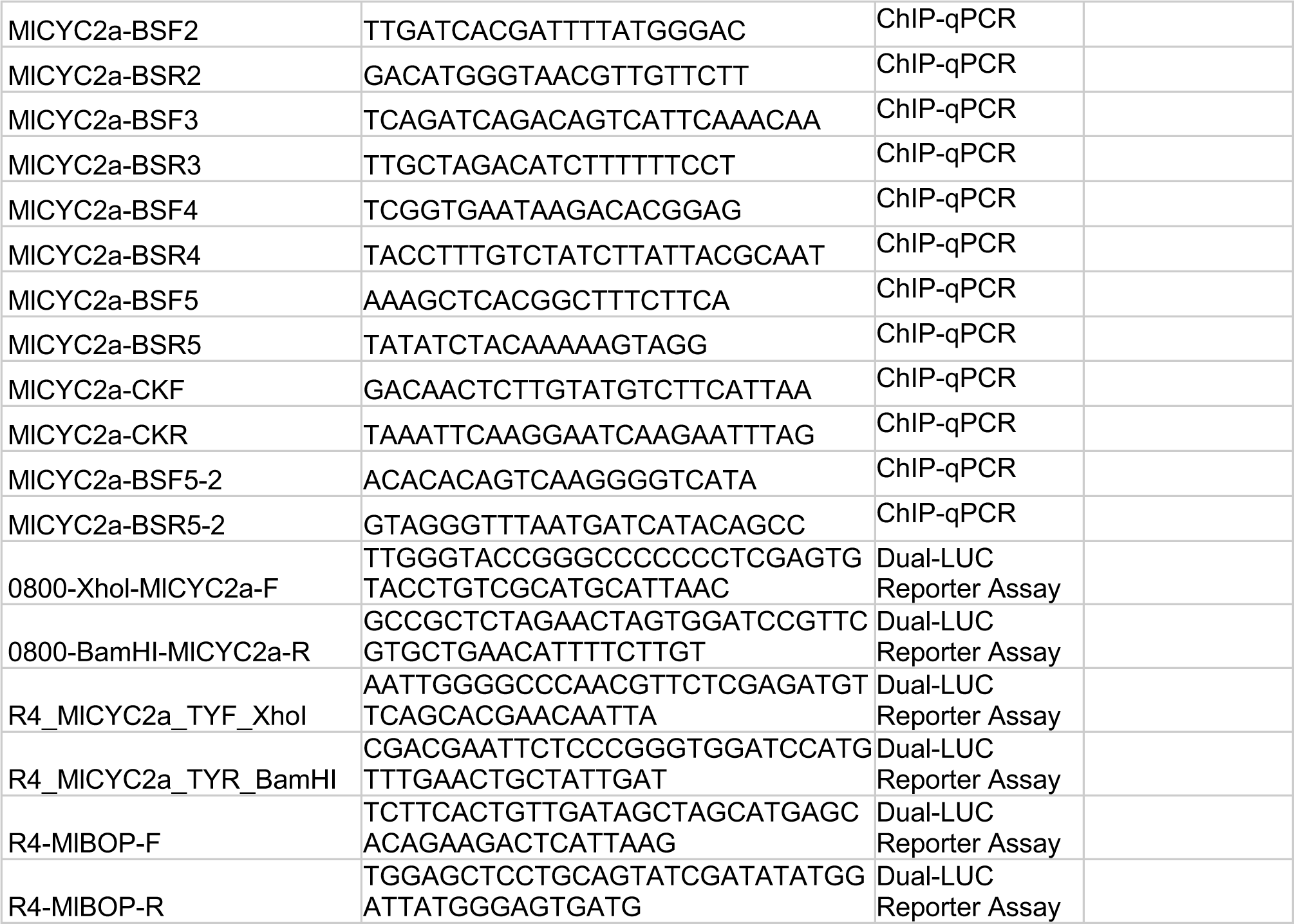
Primers used in this study.

